# Membrane Protein Modification Modulates Big and Small Extracellular Vesicle Biodistribution and Tumorigenic Potential in Breast Cancers *in vivo*

**DOI:** 10.1101/2022.09.28.510006

**Authors:** Bryan John Abel Magoling, Anthony Yan-Tang Wu, Yen-Ju Chen, Wendy Wan-Ting Wong, Steven Ting-Yu Chuo, Hsi-Chien Huang, Yun-Chieh Sung, Hsin Tzu Hsieh, Poya Huang, Kang-Zhang Lee, Kuan-Wei Huang, Ruey-Hwa Chen, Yunching Chen, Charles Pin-Kuang Lai

## Abstract

Extracellular vesicles (EVs) are released by cells to mediate intercellular communication under pathological and physiological conditions. While small EVs (sEVs; <100–200 nm, exosomes) are intensely investigated, the properties and functions of medium and large EVs (big EVs [bEVs]; >200 nm, microvesicles) are less well explored. Here, we identify bEVs and sEVs as distinct EV populations, and determine that bEVs are released in a greater bEV:sEV ratio in the aggressive human triple-negative breast cancer (TNBC) subtype. PalmGRET, bioluminescence resonance energy transfer (BRET)-based EV reporter, reveals dose-dependent EV biodistribution at non-lethal and physiological EV dosages, as compared to lipophilic fluorescent dyes. Remarkably, the bEVs and sEVs exhibit unique biodistribution profiles, et individually promote *in vivo* tumor growth in a syngeneic immunocompetent TNBC breast tumor murine model. The bEVs and sEVs share mass spectrometry (MS)- identified tumor progression-associated EV surface membrane proteins (tpEVSurfMEMs), which include SLC29A1, CD9 and CD44. tpEVSurfMEM depletion attenuates EV lung organotropism, alters biodistribution, and reduces protumorigenic potential. This study identifies distinct *in vivo* property and function of bEVs and sEVs in breast cancer, which suggest the significant role of bEVs in diseases, diagnostic and therapeutic applications.

## 1. Introduction

Extracellular vesicles (EVs) are nanovesicles with a lipid bilayer released by cells to mediate cell–cell communication[1]. Small EVs (sEVs) are <100–200-nm wide, whereas medium or large EVs (big EVs [bEVs]) are >200-nm wide[2]. We termed the bEVs and sEVs as such as their subcellular origins have not been identified, unlike microvesicles and exosomes[3]. Cells also release non-membranous nanoparticles (<50 nm) (exomeres[4] and supermeres[5]) with distinct cargo signatures. Moreover, aggressive cancers release large oncosomes (1–10 µm) that contain oncoproteins[6].

Breast cancer is a highly heterogeneous disease accompanied by a diverse set of clinical characteristics and molecular subtypes[7]. Among the breast cancer subtypes, triple-negative breast cancers (TNBC) lacking estrogen receptor (ER), progesterone receptor (PR), and human epidermal growth factor receptor 2 (HER2) exhibits the worst prognosis, followed by the HER2+, luminal B, and luminal A subtypes[8]. Cancer EVs and the delivered bioactive cargos are important in cancer progression, including tumor growth, angiogenesis, and metastasis[9]. However, whereas most breast cancer EV studies focused on sEVs, fewer investigations methodically explored bEVs in breast cancers and compared the *in vivo* properties and functions. Moreover, whether the breast cancer subtypes release a differential abundance of specific EV subtypes (bEVs and sEVs) remains largely unexplored.

Tracking EVs *in vivo* remains difficult due to their size, where EVs typically require labeling for subsequent detection, especially *in vivo*[10]. EV surface reporters function by labeling the membrane, which rely on EV surface properties to yield optimal labeling, coverage, and signal intensity.[11] Lipophilic dye-based labeling (PKH26, DiD, DiR) typically takes a short time to perform but form nanomicelles to yield false positive signals^[11a, 12]^.

Furthermore, PKH26 and DiR, initially utilized for cell tracking, have an *in vivo* half-life of 5 to >100 days[13], which may not reflect the true spatiotemporal property of the labeled EVs. In parallel, high-dose labeled EVs (∼50–100 μg) are commonly applied in mouse models^[11a, 12a,14]^, which is unlikely to correspond to a physiologically relevant EV dosage[15]. To circumvent these potential caveats, we developed a BRET (bioluminescence resonance energy transfer)- based EV reporter (PalmGRET) to enable multi-resolution imaging and sensitive biodistribution analysis of EVs^[11b]^.

As we and others have reported, surface membrane proteins can dictate EV *in vivo* dynamics.^[11b, 16]^ Specifically, proteins expressed on the EV surface (e.g. tetraspanins, galectins, and integrins) enable EVs to interact with membrane proteins on specific recipient cells *via* ligand-receptor mechanism for subsequent bioactive cargo entry and delivery.[17] Both bEVs and sEVs are found to express surface membrane proteins that correspond to their cells of origin.[18] However, the degree of expression of these membrane proteins in both bEV and sEVs and how it could affect their biodistribution remains largely uncharted, and thus warrant detailed explorations.

In the present study, we examined the bEV and sEV release index in different breast cancer subtypes, and systematically compared EV labeling modalities, dosages, and lethality.

We subsequently identified innate biodistribution, organotropic tumor progression-associated EV surface membrane proteins (tpEVSurfMEMs), and individual capacities of TNBC-derived bEVs and sEVs to induce tumorigenesis *in vivo*. These findings have important implications for bEV- and sEV-mediated cell-cell communication in disease and therapy.

## 2. Results

2.1. TNBC cells release more bEVs per and sEVs than normal cells

We isolated bEVs and sEVs from human TNBC, HER2+, and ER/PR+ luminal breast cancer cell lines using differential centrifugation and compared them to EVs from human mammary M10 epithelial cells (**Figure 1a, b**). Nanoparticle tracking analysis (NTA) revealed that the TNBC (MDA-MB-231, Hs578T, BT-549) and ER/PR+ (MCF-7) breast cancer cells released comparable amounts of bEVs and sEVs per cell, whereas HER2+ (SK-BR-3) breast cancer and M10 epithelial cells released significantly fewer bEVs than sEVs per cell (**Figure 1c, d**). Upon normalization with sEVs, all human TNBC and ER/PR+ breast cancer cells released higher amounts of bEVs compared to the M10 epithelial cells (**Figure 1e**).

**Figure 1.**
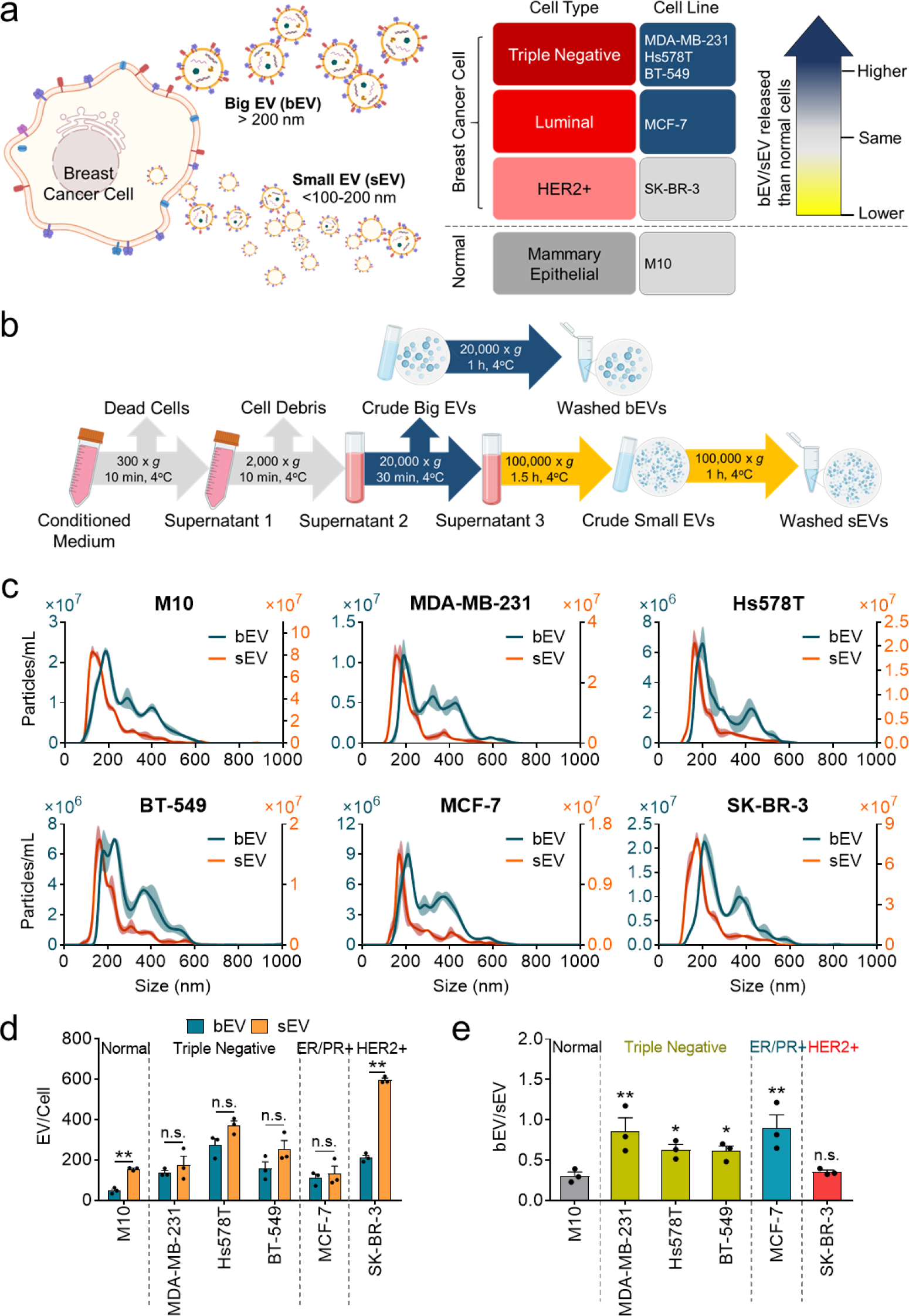
Breast cancers release more bEV per sEV than normal cells. a, Diagram showing cancer cell release of bEVs and sEVs. The highly aggressive TNBC and ER/PR+ breast cancer cells released more bEVs and sEVs as compared to normal cells. **b,** Schematic for bEV and sEV isolation using differential centrifugation. **c-e,** NTA of bEVs and sEVs isolated from human breast cancers and normal epithelial cells. (**c**) Particle size distribution plots of bEVs and sEVs isolated from normal human mammary epithelial (M10) cells, human triple-negative (MDA-MB-231, Hs578T, BT-549), ER/PR+ (MCF-7), and HER2+ (SK-BR-3) breast cancer cells. (**d**) Human TNBC and ER/PR+ breast cancers released comparable amounts of bEVs and sEVs per cell while human HER2+ breast cancer and normal human epithelial cells released significantly lower amounts of bEVs than sEVs per cell (determined from three independent experiments). (**e**) Human TNBC and ER/PR+ breast cancer cells released significantly higher bEV-to-sEV ratios as compared to HER2+ cells and normal human epithelial cells. n.s., *p* > 0.05; \**p* < 0.05; \*\**p* < 0.01 with 1-way ANOVA followed by Dunnett’s post hoc test vs. the control.

The high bEV:sEV ratio of the aggressive TNBC cells suggested that while sEVs enhanced carcinogenesis[19], bEVs may also be important during tumor progression *via* circulation and distribution *in vivo*. To investigate TNBC-secreted bEVs and sEVs, we used murine TNBC-resembling 4T1 cancer cells as a syngeneic, immunocompetent breast tumor model. Similar to human TNBC cells, the 4T1-derived bEVs (269.27 ± 11.0 nm) demonstrated significantly larger particle sizes than the sEVs (221.67 ± 7.6 nm) (**Figure 2a**), which indicated that 4T1 cells are an ideal donor of TNBC-derived bEVs and sEVs.

**Figure 2.**
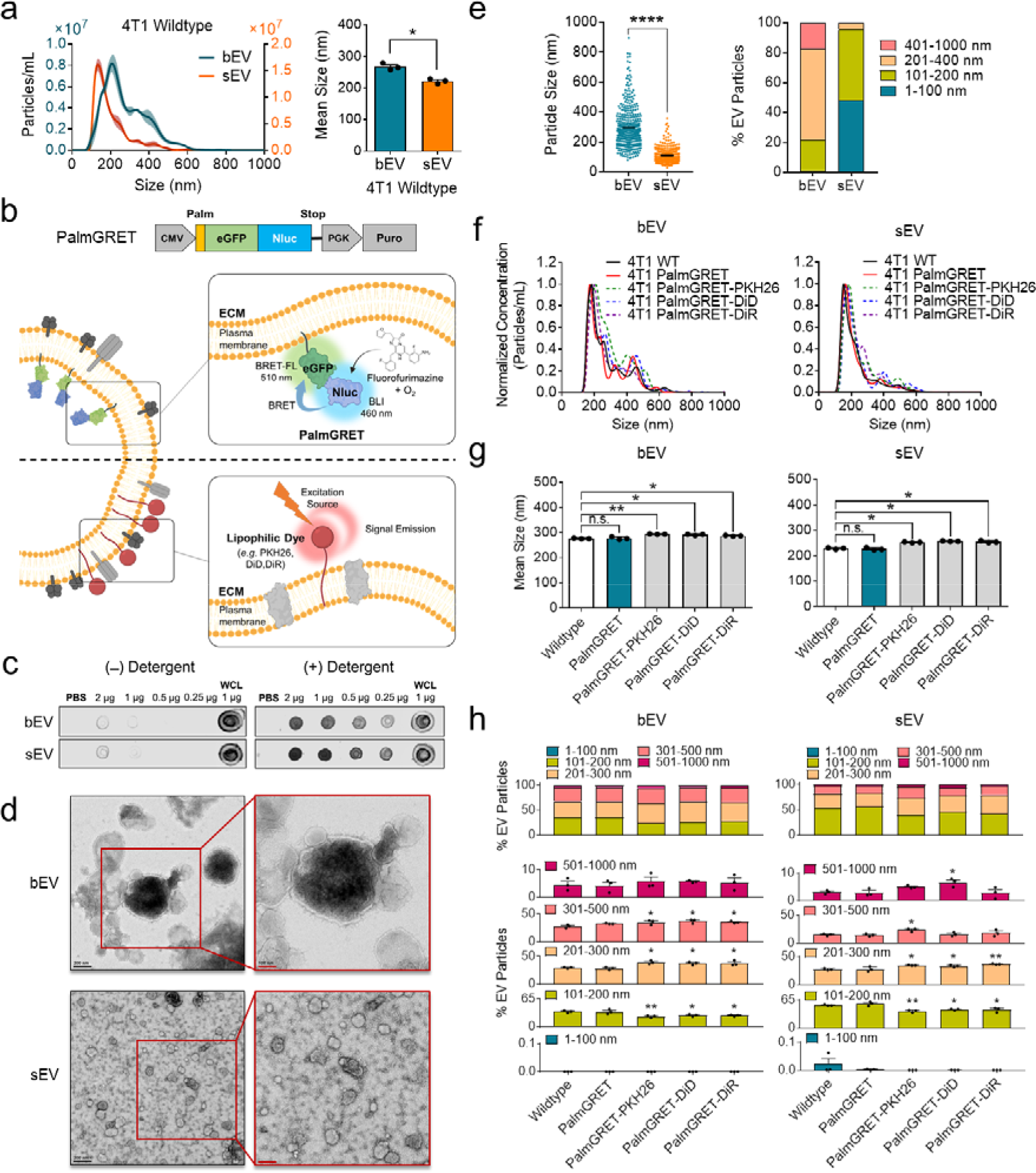
Comparison of size and detection limit between PalmGRET- and dye-labeled bEVs and sEVs a, NTA of 4T1-bEVs and -sEVs demonstrating distinct size distributions (left; representative data of three independent experiments) and mean sizes (right; determined from three independent experiments). **b,** Schematic of inner EV membrane labeling using PalmGRET (top) and outer EV membrane labeling using lipophilic fluorescent dyes (bottom). **c,** Dot blot analysis showing that PalmGRET labeled the inner membrane of bEVs and sEVs. The positive control was whole cell lysates (WCL). **d,** Representative transmission electron microscope (TEM) images of 4T1-PalmGRET bEVs and sEVs. The same imaging parameters were used to acquire bEV and sEV TEM images. Images on the right of each row depict the enlarged images of the boxed regions (red). Black bar, 200 nm; red bar, 100 nm. **e,** Mean particle size (left) and particle size distribution (right) of 4T1-PalmGRET bEVs and sEVs determined from TEM image analysis. The bEV and sEV particles (*N*=400 per group) in the captured TEM images were analyzed using Fiji software (ImageJ, NIH) to measure individual particle sizes (left). Size distribution analysis of bEVs and sEVs (right) demonstrating that bEVs are largely composed of 201–400-nm particles while sEVs mainly consist of 1–100- and 101–20-nm particles. \*\*\*\**p* < 0.0001 from 2-tailed Student’s *t*-test. **f-h,** NTA of PalmGRET- and dye-labeled 4T1-EVs. Size distribution **(f)** and mean size **(g)** of bEVs and sEVs labeled with PalmGRET or co-labeled with PKH26, DiD, and DiR are shown. Analysis of particle size composition of the bEVs and sEVs **(h)** showed that PalmGRET did not change the size distribution in all size divisions as compared to the WT EV controls. The 101–200- nm particles were decreased while the 201–300-nm and 301–500-nm particles were increased among the lipophilic dye-labelled bEVs. Similarly, the 101–200-nm particles were decreased and 201–300-nm particles were increased among the lipophilic dye-labelled sEVs. Data are from three independent experiments. n.s., *p* > 0.05; \**p* < 0.05; \*\**p* < 0.01 with 1-way ANOVA followed by Dunnett’s post hoc test vs. the control.

### 2.2. Comparison of size and detection limit between PalmGRET- and dye-labeled bEVs and sEVs

To accurately track EV spatiotemporal properties *in vivo*, it is critical that the labeled EVs resemble their unlabeled counterparts (mean size, size distribution). PalmGRET labels the inner EV membrane leaflet via its palmitoylation moiety to enable multi-resolution imaging and sensitive biodistribution analysis of EVs[20] (**Figure 2b, top**). Contrastingly, lipophilic dyes label EVs by insertion into the lipid bilayer (**Figure 2b, bottom**). To generate the 4T1-PalmGRET stable cell line, we transduced 4T1 cells with lentivirus containing PalmGRET plasmids. PalmGRET-EVs isolated from 4T1 cells stably expressing PalmGRET underwent dot blot analysis to validate the PalmGRET inner membrane labeling specificity (**Figure 2c**). If PalmGRET is located in the inner membranes, antibodies would be unable to access the reporter protein unless the EV membrane is disrupted. Conversely, the addition of Tween 20 disrupts the EV membrane to allow access to antibodies to bind the PalmGRET. A strong PalmGRET signal was detected only in the presence of 0.1% (v/v) Tween 20 in a dose-dependent manner, which indicated inner EV membrane-specific labeling of the bEVs and sEVs derived from stable 4T1-PalmGRET cells. This finding validated the EV inner membrane specificity of PalmGRET for labeling bEV and sEVs.

EVs exhibit refractive indices of 1.37–1.45 resulting in a detection limit of ≥50 nm for NTA.[21] While NTA is conventionally used to determine EV size distribution, its detection limit may not be able to account for very small EVs and could lead to a result that differs from the actual distribution. Transmission electron microscopy (TEM) analysis was performed as an orthogonal method to characterize bEVs and sEVs isolated by differential ultracentrifugation (**Figure 2d)**. TEM imaging revealed that bEVs and sEVs had mean particle sizes of 295.85 ± 6.7 nm and 110.14 ± 2.5 nm, respectively (**Figure 2e, left**). Moreover, the bEVs largely consisted of 201–1000-nm particles (78.13%) while sEVs were mainly composed of 1–200-nm particles (95.59%; **Figure 2e, right**). The 201–400-nm (61.06%) and 401–1000-nm (17.07%) particles constituted most bEVs (78.13%), whereas the 1–100-nm (48.21%) and 101–200-nm (47.38%) particles composed the majority of sEVs (95.59%). Notably, bEVs comprised 101–200-nm (20.67%) and a minute amount of 1–100- nm (1.20%) particles while sEVs contained a residual amount of 201–400-nm (4.41%) particles. Although differential centrifugation-based EV isolation could not completely separate bEVs from sEVs, the orthogonal TEM analysis provided strong evidence that the cells released different EV subpopulations with high abundances in the determined size ranges (bEVs, 201–1000 nm; sEVs, 1–200 nm).

While TEM analysis provides accurate particle size determination based on individual particle measurements, the sample preparation time is long and prevents efficient comparative study of multiple EV samples. Alternatively, NTA enables direct EV sample measurements with a short processing time, thereby allowing the analysis of a larger number of samples. Therefore, NTA of EVs isolated from wild-type 4T1 (4T1-WT, control) and stable 4T1- PalmGRET cells was performed to investigate whether PalmGRET or the lipophilic dyes would alter EV size (**Figure 2f**). Concurrently, we examined PKH26-, DiD-, or DiR -labeled 4T1-PalmGRET-EVs. The PalmGRET-bEV and -sEVs exhibited similar size distribution patterns when compared to the control. By contrast, the PKH26-, DiD-, and DiR-labeled bEVs and sEVs demonstrated increased size distributions. Similarly, the mean size of the WT- and PalmGRET-EVs did not vary, whereas the dye-labeled bEVs and sEVs exhibited significantly larger mean sizes (**Figure 2g**).

We then further compared the particle distributions between the WT-, PalmGRET-, and dye-labeled EVs at different size ranges (**Figure 2h**). The PalmGRET-bEVs and -sEVs did not vary significantly compared to the WT-EVs, which suggested that PalmGRET labeling retained the original particle size heterogeneity for both bEVs and sEVs. Conversely, the lipophilic dye labeling increased EV sizes at specific size ranges, thereby altering the overall EV size distribution patterns.

### 2.3. bEVs and sEVs exhibit differing composition sizes and densities

To determine detailed bEV and sEV size composition, freshly isolated EVs underwent sucrose density gradient fractionation followed by biophysical characterizations (**Figure 3a**). Bioluminescent assay (BLI) and NTA revealed that bEVs and sEVs were enriched in overlapping but different sucrose density fractions (**Figure 3b–d**). The BLI signals detected pre- (**Figure 3b**) and post-fraction pelleting (**Figure 3c**) positively corroborated with the particle concentration determined by NTA (**Figure 3d**). Whereas bEVs were primarily enriched in fractions 4–8, sEVs were mainly present in fractions 3–6. In accordance with these results, western blotting of the pelleted EVs demonstrated that PalmGRET coincided with EV markers (CD81 and Alix [ALG-2-interacting protein X]) in the bEV and sEV enriched fractions (**Figure 3e**). Alix was more enriched in the sEV than bEV enriched fractions. The total particle composition analysis revealed that the bEVs were composed of a higher percentage of 201–400- and 401– 1000-nm particles as compared to the sEVs (**Figure 3f**). Additional characterization to compare the particle composition profiles in each density fraction demonstrated that the bEVs and sEVs had significantly different percent particle size distributions in the 101–200-, 201– 400-, and 401–1000-nm ranges (**Figure 3g**).

**Figure 3.**
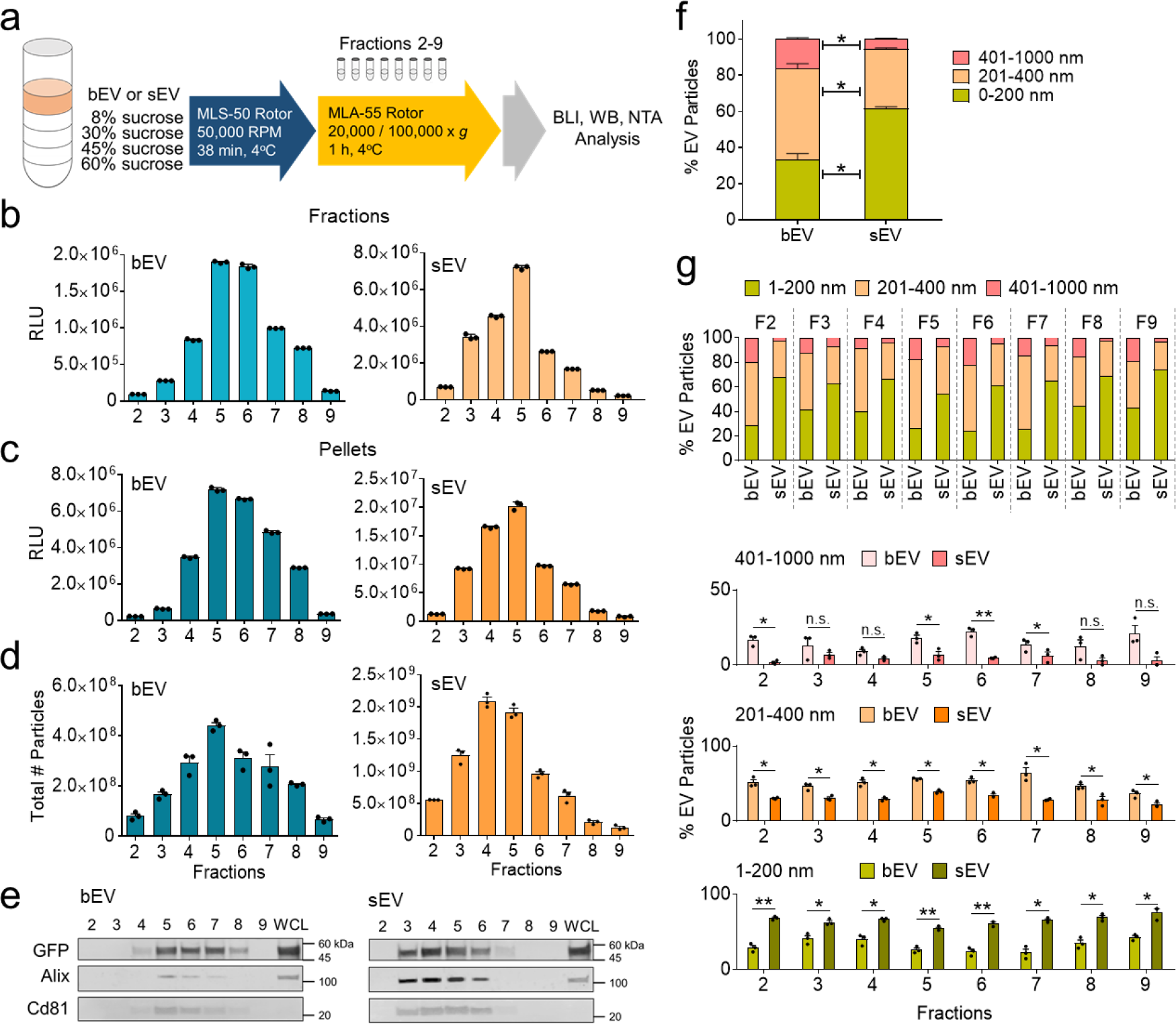
bEVs and sEVs differ in size composition and density. a, Flowchart of sucrose gradient fractionation of 4T1-PalmGRET-bEVs and -sEVs. The sucrose density gradient fractions 2 (1.054 g-cm^-3^), 3 (1.086 g-cm^-3^), 4 (1.117 g-cm^-^3), 5 (1.139 g-cm^-3^), 6 (1.163 g-cm^-3^), 7 (1.187 g-cm^-3^), 8 (1.208 g-cm^-3^), and 9 (1.219 g-cm^-3^) were collected for subsequent analyses. **b, c,** Nluc assay of the EVs following fractionation **(b)** and fraction pelleting **(c)**. **d,** NTA of pelleted sucrose gradient fractions. **e,** Western blotting of bEV (left) and sEV (right) proteins from the pelleted fractions revealing that PalmGRET (49 kDa) coincided with the Alix (96 kDa) and CD81 (26 kDa) EV markers. The positive control was WCL. **f,** Total particle size composition analysis (fractions 2–9) showing that bEVs were composed of a significantly higher percentage of 201–400-nm particles (bEV: 50.47% vs. sEV: 32.85%) and 401–1000-nm particles (bEV: 16.03% vs. sEV: 5.42%), whereas sEVs were primarily comprised of 101–200-nm particles (bEV: 33.50% vs. sEV: 61.73%). **g,** Particle analysis of sucrose gradient fractions demonstrating that bEVs and sEVs had significantly different size distribution in the 101–200-nm (all fractions), 201–400-nm (all fractions), and 401–1000-nm (fractions 2, 5–7) size ranges. n.s., *p* > 0.05; \**p* < 0.05; \*\**p* < 0.01 with 2-tailed Student’s *t*- test.

These results demonstrated that the bEVs and sEVs consisted of particles with lower and higher densities, respectively. Furthermore, while the density fractionation demonstrated that the bEVs and sEVs shared overlapping densities, the fractions comprised significantly different particle size composition profiles.

### 2.4. bEVs have a greater surface area per particle than sEVs

Fluorofurimazine (FFz) is a modified version of furimazine (Fz) with improved water solubility and hence greater reactivity with Nluc luciferase (enhanced signal)[22]. We tested the applicability of FFz, and demonstrated that FFz could be applied to enhance BLI and BRET-induced fluorescence activities of PalmGRET-labeled bEVs and sEVs (**Figure S1, S2**).

We examined the BLI properties of PalmGRET-bEVs and -sEVs by measuring BLI and BRET-FL activities supplemented with FFz over time. Following normalization to EV particles, Nluc (**Figure 4a**) and BRET-FL (**Figure 4b**) demonstrated that bEVs exhibited higher signal intensity per particle when compared to the sEVs. As sEVs are smaller (**Figure 2a, e**), they should be of lower volume than bEVs. Furthermore, as PalmGRET labels the EV membrane (but not EV cytosol), a fixed amount of protein should therefore contain a greater number of sEVs than bEVs, thereby yielding a higher PalmGRET signal per protein over time in sEVs. Indeed, Nluc (**Figure 4c**) and BRET-FL (**Figure 4d**) per protein (ng) were higher in the sEVs than bEVs, demonstrating that sEVs are of lower volume than bEVs. The half-life of the signals and BRET ratios (**Figure 4e**) were comparable under both normalization settings, which indicated that PalmGRET functioned similarly in the labeled bEVs and sEVs. The protein concentration assay indicated no difference in the total protein amount between the bEVs and sEVs from each EV isolation batch (**Figure 4f**). Normalizing the detected PalmGRET signal against the amount of protein at a single time point (**Figure 4g**) revealed significantly lower relative light units (RLU) per protein (ng) in the bEVs than sEVs, again validating the finding that bEVs are of greater volume than sEVs. As bEVs are greater in volume and hence surface area, they should contain more protein and EV membrane-labeling PalmGRET, respectively. Indeed, bEVs contained a significantly higher amount of protein and reporter signal per particle (∼2-fold) when compared to sEVs (**Figure 4h, i**). Contrastingly, the difference in PalmGRET signal between the bEVs and sEVs was less (∼1/5-fold) when the signal was normalized against the protein amount (**Figure 4g**). Therefore, to minimize reporter signal differences between bEVs and sEVs as a result of the varying sizes and volumes (**Figure 4g–i**), we applied EVs in protein amounts (instead of particle numbers) in subsequent experiments.

**Figure 4.**
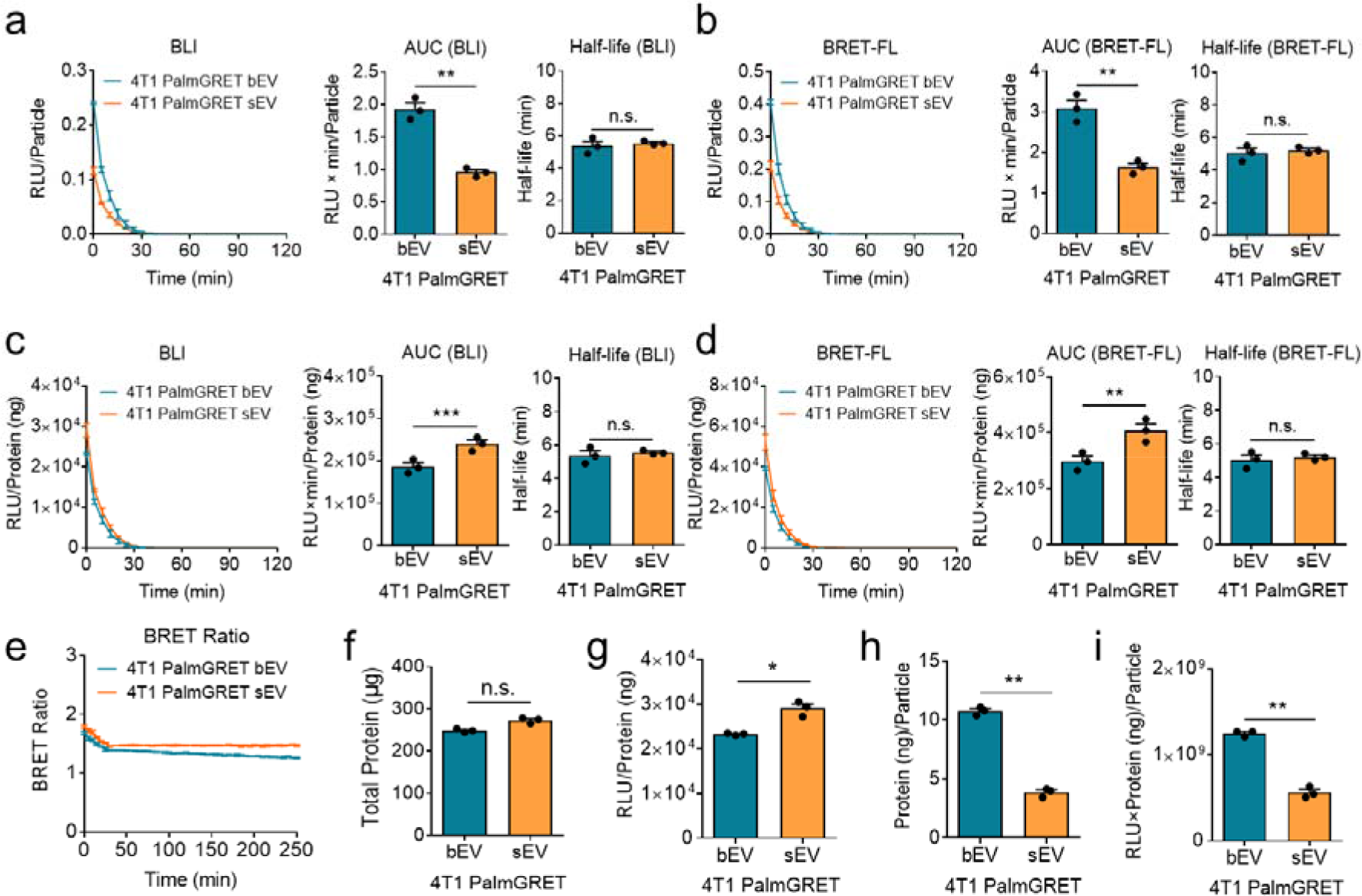
PalmGRET demonstrates that bEVs contain more protein per particle than sEVs. a, b, Following normalization to EV particles, Nluc **(a)** and BRET-FL **(b)** demonstrated that bEVs had higher signal intensity (area under the curve [AUC]) per particle when compared to sEVs. **c, d,** Following normalization to EV protein amount, Nluc **(c)** and BRET-FL **(d)** revealed that sEVs had higher RLU (relative light unit) per protein (ng). **e,** Half-life of the signals and BRET ratio were comparable under both normalization settings, which indicated that the PalmGRET reporter functioned similarly in bEVs and sEVs. **f,** Protein concentration assay indicated no significant difference in total protein amount between bEVs and sEVs from each EV isolation batch. **g,** sEVs exhibited higher RLU per protein (ng) than bEVs. **h,** Bar graphs showing significantly more protein (ng) per bEV particle than sEVs. **i,** Bar graphs indicating bEVs exhibited higher RLU × ng/particle.

### 2.5. PalmGRET labeling exhibits a high dynamic range (HDR), and enables sensitive EV biodistribution analysis at non-lethal low dosages *in vivo*

We compared the detection limits and linearity between the PalmGRET and DiR signals of 4T1-PalmGRET-DiR-bEVs and -sEVs (**Figure 5a, b**). PalmGRET signals were detected with a high correlation to EV protein concentration in bEVs (*R*^2^ = 0.9788) and sEVs (*R*^2^ = 0.9720) (**Figure 5a**). Contrastingly, DiR signals demonstrated a lower correlation in bEVs (*R*^2^ = 0.9318) and sEVs (*R*^2^ = 0.8003) and were barely at concentrations < 125 or 0–1000 ng, respectively (**Figure 5b**).

**Figure 5.**
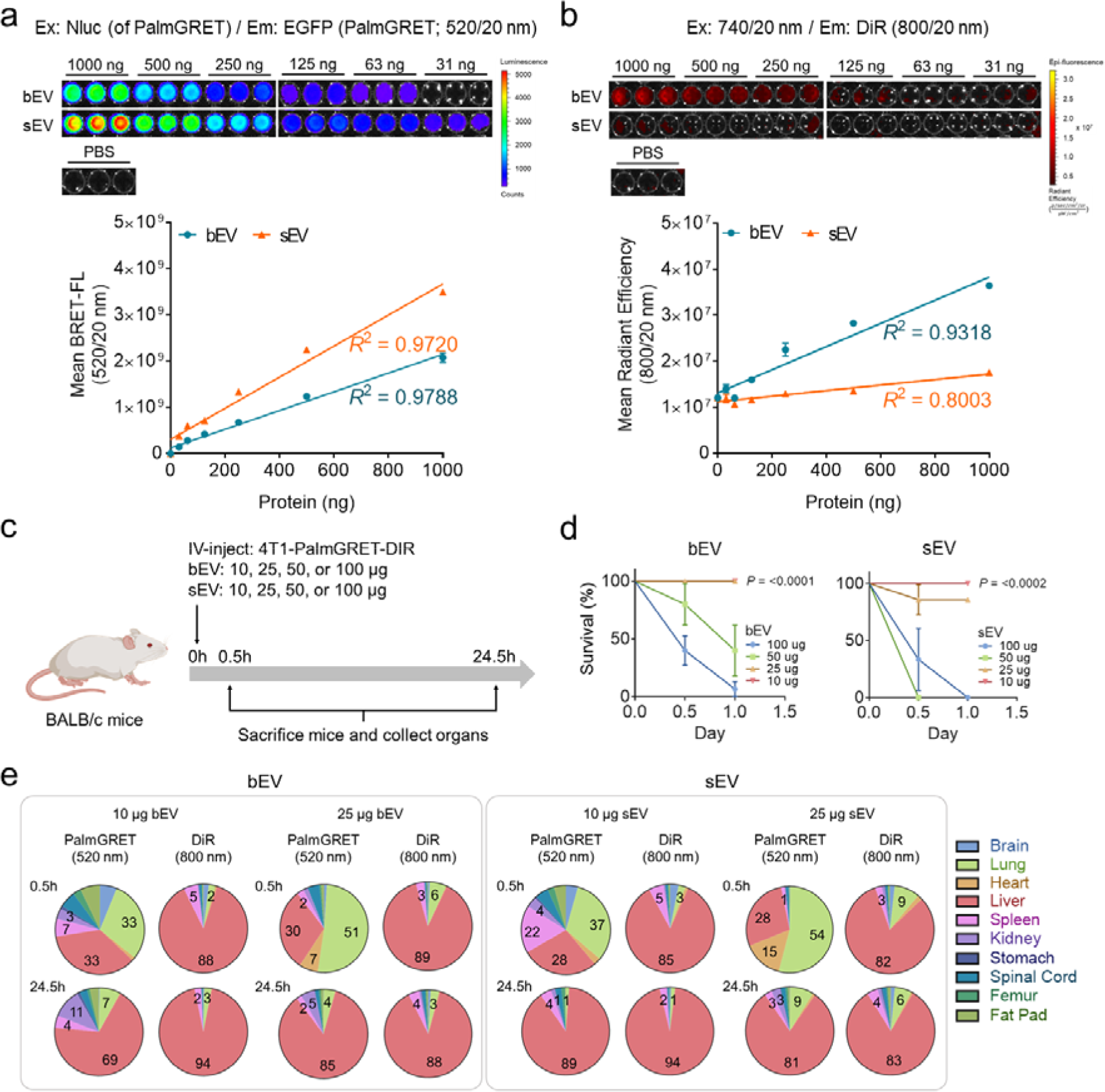
EV PalmGRET labeling enables sensitive EV biodistribution analysis at non- lethal low dosages *in vivo*. a, b, BRET-based EGFP signal of PalmGRET-DiR 4T1-EVs (**a**) demonstrating a HDR (10^9^) of PalmGRET BRET-FL signals. External excitation-based DiR signal of PalmGRET-DiR 4T1-EVs (**b**) showing a low dynamic range of DiR signals (∼10^7^). **c,** Pharmacokinetics study of 10–100 µg PalmGRET-DiR 4T1-EVs in mice. **d,** Survival rate of IV-injected mice. Only 10 µg (bEVs and sEVs) and 25 µg (bEVs only) yielded full survival among the mice (4 out of 4 per group). **e,** Differential biodistribution pattern between PalmGRET and DiR signals of IV-injected 4T1-PalmGRET-DiR EVs over time. EV distributions are presented as the percentages of the total respective signal detected.

These findings indicated that PalmGRET exhibited HDR and linearity even at low EV protein concentrations as compared to DiR.

To investigate the possible effect of dosage and DiR labeling on EV biodistribution, we administered 10, 25, 50, or 100 µg 4T1-PalmGRET-DiR-bEVs or -sEVs intravenously (IV) into immunocompetent BALB/c mice (**Figure 5c**). Unexpectedly, a high lethality rate was observed in both the 50 µg and 100 µg bEV and sEV groups, with no surviving animals at 24 h (**Figure 5d**). By contrast, there were minimal or no deaths in the 10 µg and 25 µg bEV and sEV groups at 24 h post-injection. This result indicated that ≥50 µg labeled 4T1 EVs presented a high lethality risk to the mice. We previously IV-administered 100 µg HEK293T-PalmGRET-derived EPs to mice and did not observe death within 24 h after EP injection.^[11b]^

Hence, it should be noted that the lethality risk may be dependent on the donor cell type and EV dose, as in the case of 4T1-bEVs and -sEVs.

We subsequently performed biodistribution analyses based on BRET-FL (PalmGRET) and FL (DiR) signals of the 10 µg and 25 µg bEV and sEV groups (**Figure 5e**). Acknowledging the EV size-dependent variation in the reporter signals between bEVs and sEVs (**Figure 4g**), the bEV and sEV organ biodistribution analyses are presented as percentages of the total respective signal detected (**Figure 5e**). This enabled a systemic approach to visualize the organotropism and biodistribution of bEVs and sEVs independently from each other, thereby mitigating possible bias from the signal differences between the bEVs and sEVs.

In the 25 µg groups, PalmGRET signals of both bEV and sEV exhibited the highest initial (0.5 h) distribution to the lungs followed by the liver. At 24.5 h, both bEVs and sEVs were decreased in the lungs and predominantly accumulated in the liver. In the 10 µg groups, bEVs demonstrated the highest initial (0.5 h) distribution in the lungs and liver, whereas sEVs were detected primarily in the lungs, liver, and spleen. At 24.5 h, both bEVs and sEVs were decreased in the lungs and predominantly accumulated in the liver. Oppositely, the bEV and sEV DiR signals in both the 10 µg and 25 µg groups demonstrated high distribution to the liver at 0.5 h and 24.5 h, revealing a distinct EV biodistribution pattern compared to that of PalmGRET (**Supplementary Table 3**). PalmGRET signal revealed that 4T1-bEVs and -sEVs primarily targeted the lung, liver, and spleen (sEV only) at a low dose. Notably, the temporal change in bEV biodistribution was not as prominent as that of sEVs in the 10 µg groups, which suggested that sEVs are more readily trafficked to the liver from other major organs over time, which could only be observed with low-dose EV (10 µg). The DiR signal intensity was weak and with a high background in the 10 µg and 25 µg groups (**Figure S3, Table S1, S2**). This rendered DiR signals mostly detectable in a large major organ (the liver) but not in the other smaller major organs.

### 2.6. 4T1 bEVs and sEVs share tumorigenic potential-associated EV surface membrane proteins (tpEVSurfMEM)

To investigate the proteins that may contribute to 4T1 EV lung tropism, 4T1-bEV and -sEV protein were screened based on their prior association with breast cancer tumorigenesis and metastasis to identify three EV lung tropism-related tpEVSurfMEMs: solute carrier family 29 member 1 (Slc29a1), tetraspanin Cd9, and cell surface glycoprotein Cd44 (**Figure 6a, b, Figure S4**). We next explored the roles of the identified tpEVSurfMEM candidates in directing bEV and sEV innate organotropism *in vivo* using 4T1-PalmGRET. We generated stable cell lines to knockdown (KD) the candidates of the bEVs and sEVs by transducing 4T1-PalmGRET cells using lentivirus encoding short hairpin RNAs (shRNAs) targeting a null sequence (Scramble) or the tpEVSurfMEMs **(Figure 6c, Figure S5, Table S3**).

**Figure 6.**
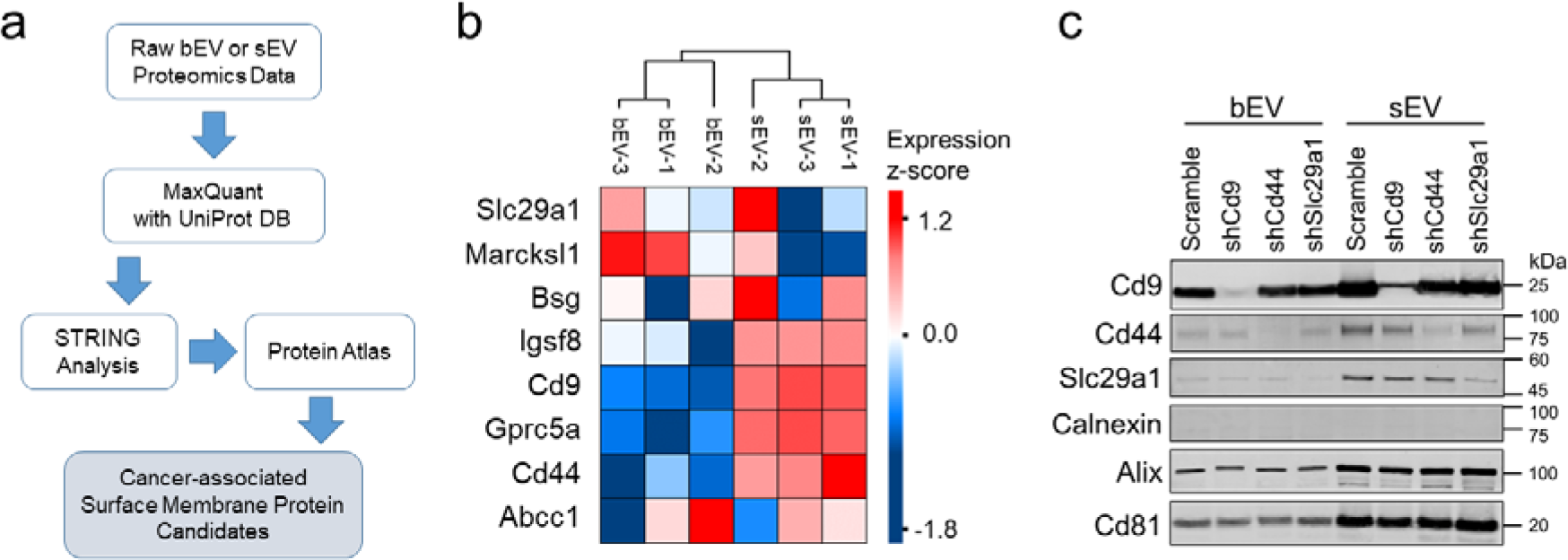
Mass spectrometry-based identification of tumorigenic potential-associated EV surface membrane proteins (tpEVSurfMEM). a, Flowchart of proteomics analysis to identify cancer-associated EV membrane protein candidates. 4T1-bEV and -sEV protein samples underwent liquid chromatography–tandem mass spectrometry (LC-MS/MS) analysis. The relative protein abundance in each sample group was determined using label-free quantification. Three biologically independent samples of bEVs and sEVs were analyzed using LC-MS/MS and MaxQuant, followed by manual STRING and the Human Protein Atlas database annotations to screen for cancer-associated membrane proteins. **b,** Heatmap illustrating the relative abundance of membrane and membrane-associated proteins identified in 4T1-PalmGRET-bEVs and -sEVs with prior cancer relation annotated in UniProt, i.e., tpEVSurfMEMs. **c)** Western blot demonstrating knockdown (KD) of selected candidates in bEVs and sEVs. 4T1-PalmGRET cells were stably transduced with lentivirus encoding shRNA targeting the null sequence (Scramble) or individual tpEVSurfMEMs followed by EV isolation and western blotting analysis: CD9 (25 kDa); CD44 (85 kDa), and SLC29A1 (55 kDa). The cell marker (negative EV control) was calnexin (85 kDa). Alix (96 kDa) and CD81 (26 kDa) were immunoprobed as EV markers.

### 2.7. Reduced tpEVSurfMEM expression attenuates lung tropism and alters bEV and sEV biodistribution

The shCd44- and shSlc29a1-EVs demonstrated significantly reduced distribution to the lungs at 0.5 h post-injection as compared to the Scramble controls, which suggested a role of Cd44 and Slc29a1 in the immediate direction of bEVs and sEVs to the lungs (**Figure 7a, e, Figure S6, Table S4)**. By contrast, the shCd9-EVs demonstrated reduced lung distribution only at 72.5 h, which indicated that Cd9 may also facilitate EV lung tropism but *via* a slower mode of action than Cd44 and Slc29a1. Significantly reduced RLU intensities of shCd9-, shCd44-, and shSlc29a1-bEVs and shCd9-sEVs were detected in the liver at 0.5 h (**Figure 7b**). Significantly reduced RLU intensities of shCd9-, shCd44-, and shSlc29a1-sEVs were detected in the spleen at 0.5 h and 72.5 h, this trend was not observed in the bEVs, which suggested that bEVs and sEVs may follow different routes toward splenic uptake. The shCd44-bEV and -sEV signals were higher in the blood at 72.5 h (**Figure 7c)**, which suggested that the reduced uptake by the major organs freed and subsequently increased circulating EV concentrations in the blood. Overall, the results demonstrated that KD expression of bEV- and sEV-shared tpEVSurfMEMs reduced lung tropism and elevated circulating EV levels in the blood.

**Figure 7.**
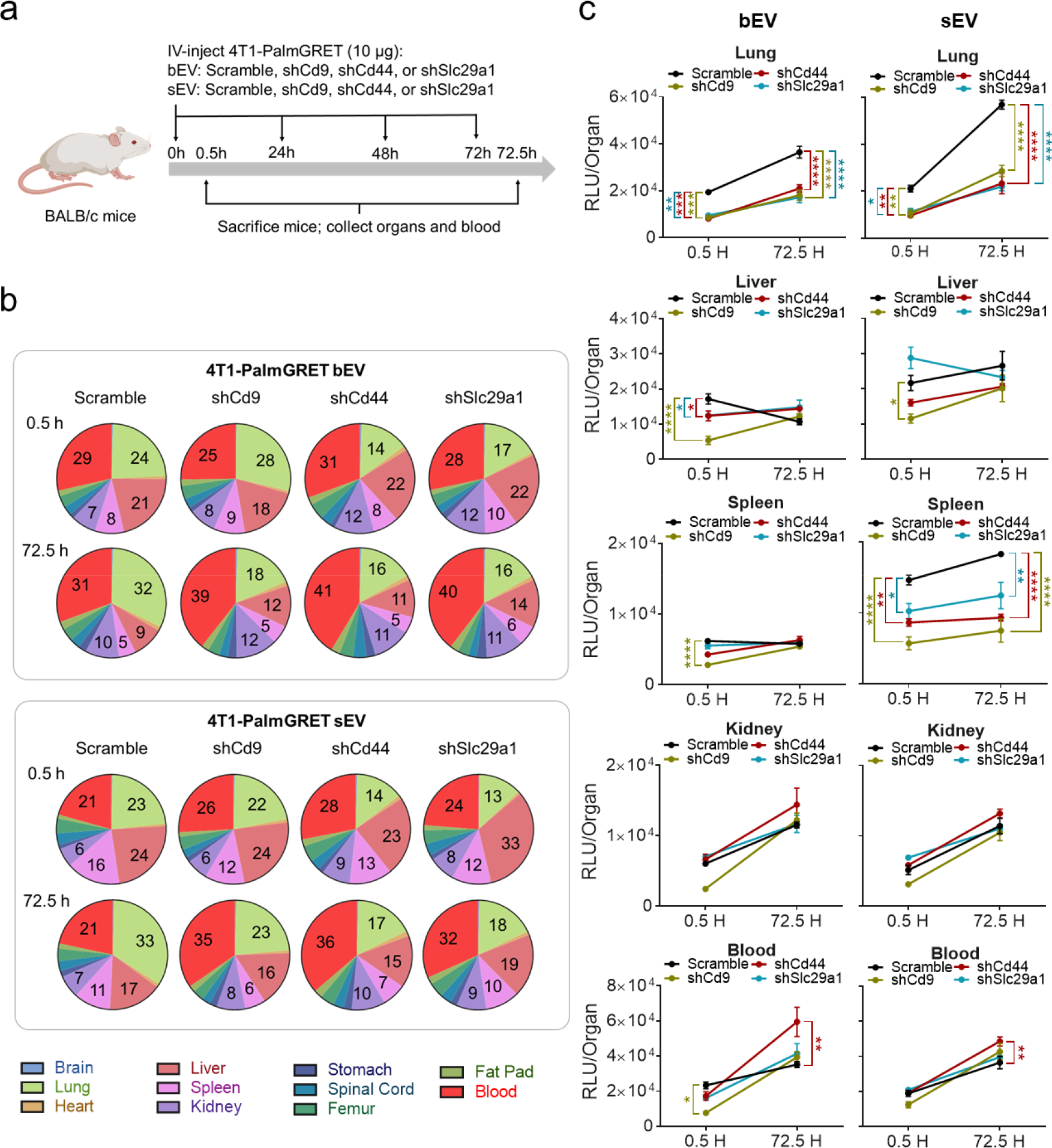
Reduced tpEVSurfMEM expression attenuates lung tropism and alters bEV and sEV biodistribution. a, Schematic for the redirection of EV organotropism where 4T1 BALB/c mice were IV-injected with EVs (10 µg) at 0, 24, 48, and 72 h through the tail vein, and the organs were collected for biodistribution analysis at 0.5 h and 72.5 h. **b,** EV biodistribution percentage pie chart demonstrating reduced lung tropism and altered distribution at organ system level of shCd9-, shCd44-, and shSlc29a1-4T1-PalmGRET-sEVs and -bEVs when compared to the Scramble controls over 3-day EV treatment. shCd44- and shSlc29a1-EV lung distribution was strongly decreased beginning at 0.5 h, which persisted to 72.5 h. shCd9-EVs demonstrated reduced lung distribution only at 72.5 h. The EV distribution percentage for each organ at different time points were calculated as the mean organ RLU per mean RLU of all organs. *N* = 3 mice per group with technical triplicates for each sample. **c,** Total EV signals of shCd9-, shCd44-, and shSlc29a1-4T1-PalmGRET-sEV and bEV distribution in the lung, liver, kidney, spleen, and blood compared to that of the Scramble controls. Data are the mean ± SEM. \**p* < 0.05; \*\**p* < 0.01 with 1-way ANOVA followed by Dunnett’s post hoc test vs. Scramble controls.

### 2.8. bEVs and sEVs individually promote breast cancer tumor growth

To mimic and investigate if constant exposure to cancer bEVs and sEVs in the circulation had a protumorigenic function, syngeneic immunocompetent 4T1 tumor-bearing mice received IV 10 µg 4T1-PalmGRET-Scramble- and -shCd44-bEVs and -sEVs, or PBS (control) three times weekly (**Figure 8a**). Scramble-bEV and -sEV groups yielded significantly higher tumor weights and volume, as well as lower body weights as compared to the control, which suggested that bEVs and sEVs individually induced a protumorigenic effect (**Figure 8b, c**). Moreover, the shCd44-bEV and -sEV groups demonstrated similar tumor weights and volumes as compared to the control, which indicated that depleting Cd44 in the EVs attenuated the protumorigenic potential of 4T1-EVs (**Figure 8b**). Biodistribution analysis of the EVs in the tumor-bearing mice revealed distinct distribution profiles between the Scramble- and shCd44-EVs, where shCd44-sEVs and -bEVs significantly reduced lung distribution and increased circulation in the blood (**Figure 8d, Figure S7, Table S5**). At day 14 and 18 post-EV treatment, fewer shCd44-bEVs and -sEVs were distributed to the lungs as compared to the Scramble controls, corroborating our findings that Cd44 depletion in EVs attenuated lung tropism (**Figure 5e**). In the spleen, Scramble-sEV biodistribution was higher than that of the Scramble-bEVs. While there was no difference between shCd44-bEV and Scramble-bEV biodistribution in the spleen, significantly reduced shCd44-sEV levels were detected in the spleen compared to Scramble-sEVs, which suggested that Cd44 depletion in the tumor-bearing mice might have reduced the innate delivery of 4T1-sEVs to the spleen. There were higher levels of both shCd44-bEVs and -sEVs in the blood as compared to the Scramble controls, which indicated that the reduced EV uptake by the major organs consequently increased shCd44-EV circulation in the blood. Both shCd44-bEVs and -sEVs exhibited significantly decreased distributions to the tumor as compared to the Scramble controls. Furthermore, both shCd44-bEVs and -sEVs exhibited significantly decreased distributions to the tumor as compared to the Scramble controls (**Figure 8e**). We validated bEV and sEV distribution in the lung, spleen, and tumors *via* super-resolution microscopy (**Figure 8f, g, Figure S8a**). Subcellular EV signals were uniformly distributed in the spleen and tumor, while EV distribution in the lungs was sporadic. (**Figure 6f**) Z-stack imaging demonstrated intracellular bEV and sEV uptake in the spleen, lung, and tumors of mice treated with 4T1-PalmGRET-Scramble- or shCd44-bEVs or -sEVs (**Figure S8b**). Altogether, our results reveal that Cd44 is pivotal in promoting tumor growth individually *via* either bEVs or sEVs and that reduced Cd44 expression redirected organotropism and subsequent change in biodistribution at organ system level.

**Figure 8.**
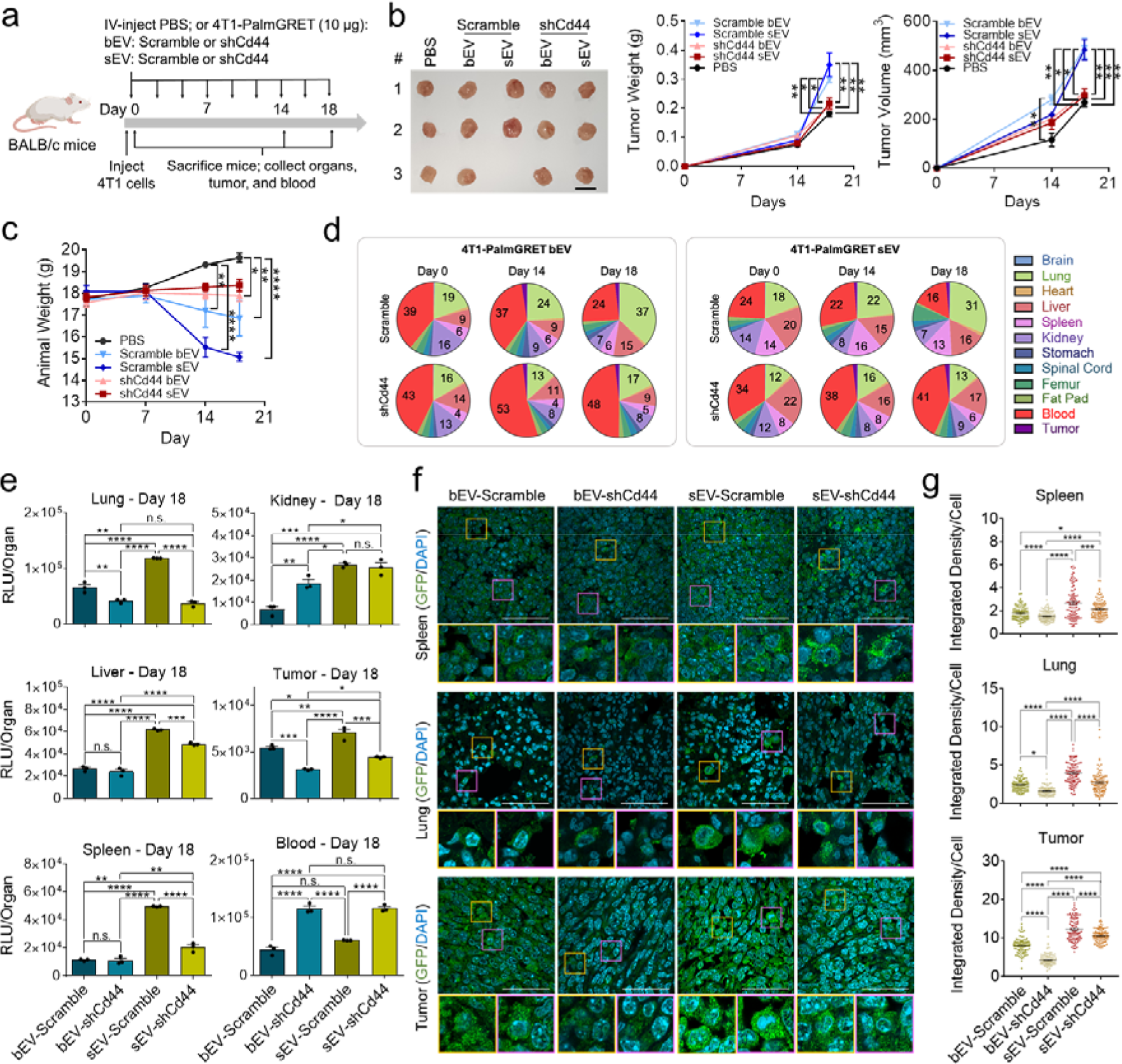
Lung-tropic cancer-derived EVs promote breast cancer tumor growth; Depleting CD44 in cancer bEVs and sEVs reduces EV cellular uptake in the spleen, lung, and tumor. a, Schematic of tracking biodistribution and tumorigenic effect of IV lung-tropic 4T1-EVs in the syngeneic 4T1-BALB/c orthotopic breast cancer model. BALB/c mice were subcutaneously injected with 5 × 10^4^ 4T1 cells in the fat pad. The mice received PBS, 4T1- PalmGRET-Scramble bEV/sEV (10 µg), or -shCd44 bEV/sEV (10 µg) thrice per week for 18 days. The blood and organs were harvested at day 0, 14, and 18 for biodistribution analysis. **b,** Isolated orthotopic 4T1 tumors (left) on day 18. Bar, 10 mm. Tumor weight (middle) and tumor volume (right) at day 14 and 18. *N* = 3 per group. \**p* < 0.05; \*\**p* < 0.01; with 1-way ANOVA followed by multiple group comparisons. **c,** Animal weight from day 0 to 18 post- 4T1 implantation and EV administration. *N* = 4 mice per group. \**p* < 0.05; \*\**p* < 0.01; \*\*\*\**p* < 0.0001 with 1-way ANOVA followed by multiple group comparisons. **d,** EV biodistribution percentage pie charts showing significantly reduced lung distribution and increased circulation in the blood of shCd44-sEVs and -bEVs in the tumor-bearing mice. The proportions for each organ at different time points were calculated as the mean organ RLU per mean RLU of all organs. *N* = 3 mice per group with technical triplicates. **e,** Bar graphs comparing the Nluc activity among the IV-injected 4T1-PalmGRET-shCd44-bEVs and -sEVs (after 18-day EV treatment) as compared with the Scramble controls are shown. \**p* < 0.05; \*\**p* < 0.01 1-way ANOVA followed by multiple group comparisons. **f, g,** Super-resolution imaging **(f)** and quantification of EV signals **(g)** in the spleen, lung, and tumor sections of mice injected with 4T1-PalmGRET-Scramble/shCd44-bEVs or -sEVs. **f,** Representative super-resolution confocal images of the spleen, lung, and tumor sections collected after 18- day treatment with 4T1-PalmGRET-Scramble or -shCd44-bEVs or -sEVs. All tissue section groups were prepared under similar conditions. The nuclei were stained with DAPI and 4T1- PalmGRET was immunoprobed by anti-GFP antibody followed by Alexa Fluor 568- conjugated secondary antibody. Bottom inset of each row depicts the enlarged images of the boxed regions (yellow and purple). Bar, 50 µm. **g,** Quantification of EV signals in the tissue sections in (**f**) demonstrated significantly decreased integrated density per cell in the lung and tumor of mice treated with shCd44-bEVs and -sEVs as compared to the Scramble controls. Only the spleen of shCd44-sEV-treated mice exhibited significantly decreased integrated density per cell as compared to the shCd44-bEV and Scramble control groups. All data are the mean ± SEM. \**p* < 0.05; \*\**p* < 0.01; \*\*\**p* < 0.001; \*\*\*\**p* < 0.0001 with 1-way ANOVA followed by multiple group comparisons.

## 3. Discussion

bEVs and sEVs theoretically demonstrate significant bioactive cargo packaging capability and surface presentation, respectively [23], to facilitate intercellular communication. Hence, we speculated that both bEVs and sEVs circulate in organisms to exert biological functions. Here, we compared the biophysical properties, biodistribution, and biological effects of bEVs and sEVs from different breast cancer molecular subtypes.

Despite the seemingly overlapping fractions between the bEVs and sEVs, comparison of the particles recovered in each sucrose fraction determined that the bEVs were composed of a significantly lower percentage of 101–200 nm particles and a higher percentage of 201– 400-nm and 401–1000-nm particles when compared to the sEVs. Notably, the established NTA measurement parameters enabled the detection of a broad particle size range for a satisfactory estimate of the size distribution but may not have accurately resolved extreme size ranges such as the 401–1000-nm division[24], which was not detected in the sEVs by TEM analysis. The sEVs had higher BL activity per protein (ng) than the bEVs. Altogether, these results suggested that the bEVs and sEVs are different sized EV populations.

Subsequently, we identified PalmGRET as the optimal labeling method for *in vivo* study of bEVs and sEVs that would enable sensitive and accurate EV detection. EVs are labelled with lipophilic dyes initially designed and used to label cells for long-term observation (days to months), which yield false positive signals (micelle formation) and inaccurate spatiotemporal properties (extended half-life)^[10a, 25]^. Detailed NTA revealed that PKH26, DiD, and DiR labeling but not PalmGRET labeling altered particle size distribution and therefore the mean bEV and sEV sizes, which might have affected their spatiotemporal properties *in vivo* as previously reported for synthetic nanoparticles[26]. In fact, several studies reported that nanoparticle size could affect its uptake, biodistribution, and circulation half-life. Decreased nanoparticle and EV uptake by cells was consistently observed with increasing nanoparticle size, potentially due to the higher energy requirement for the uptake of bigger nanoparticles[27]. Larger EVs accumulated more frequently in the spleen and liver and demonstrated a shorter *in vivo* circulation half-life, possibly resulting from the triggered immune response[28]. Our biodistribution analysis showed that DiR affected the normal biodistribution of the labeled EVs over time, possibly with the prolonged half-life and lipophilic property of the dye to be recycled into cellular membranes^[10a, 29]^. The almost full lethality of high-dose EVs indicated that ≥50 μg is not a physiologically relevant dosage, and warrants attention in future investigations. Whether the lethality is attributed to the EV donor cell type (some cell-derived EVs tend to aggregate more than others; unpublished observation), the dyes, or both is unknown. Reporter genes such as PalmGRET require cellular expression for EV labeling, whereas dyes can be applied to directly stain prepared EVs^[10a]^. Therefore, the EV dose, donor cell type, and labeling method should be carefully designed and evaluated to perform optimal *in vivo* EV studies. Accordingly, we used PalmGRET and low-dose EVs (10, 25 μg) in subsequent bEV and sEV experiments.

PalmGRET demonstrated a size-, dose-, and time-dependent EV biodistribution profile. EV size-dependent distribution was observed in the 25 µg set, where more sEVs were trafficked to the heart than bEVs. More sEVs accumulated to the spleen in the 10 µg group, which supported the findings of Wen and colleagues[30]. The bEVs did not accumulate in the spleen as much as the sEVs, which suggested that an EV subtype-dependent mechanism could regulate splenic uptake. Furthermore, at 24 h-post injection, sEV signals were detected predominantly in the liver while bEV signals remained detectable in the liver, kidney, lung, and spleen, which indicated that the organs might process bEVs and sEVs differently. Our findings indicated that bEVs and sEVs are distinct EV populations which can be well circulated with differential biodistribution properties.

EV surface proteins (tetraspanins, galectins, integrins) can dictate EV *in vivo* dynamics *via* a ligand–receptor mechanism for subsequent bioactive cargo entry and delivery^[11b, 16, 17b-d]^. Here, we identified and subsequently validated the three tpEVSurfMEMs (CD44, CD9, SLC29A1) shared by bEV and sEV in directing their organotropism. As CD44 was overexpressed in invasive breast cancers and metastasis[31], we depleted Cd44 in the bEVs and sEVs and observed a significant decreased tumor delivery and growth while exerting EV population-specific change in the biodistribution pattern. These detailed observations suggested that bEVs and sEVs actively promote breast tumor growth *in vivo via* the circulation and systemic organ and tumor uptake while acting simultaneously under distinct biodistribution patterns. As the TNBC and ER/PR+ breast cancer cells released both EV subpopulations concurrently, future investigations should elucidate a synergistic (or separate) interplay between bEVs and sEVs.

## 4. Conclusion

Overall, beyond the general EV perspective, our work provided firm evidence on the equally significant role of bEVs in understanding the systemic importance of EVs *in vivo* (**Figure 9**). As both bEVs and sEVs interplay during disease progression, it is crucial to elucidate their biodistribution profiles to understand how they might potentially mitigate disease development. Furthermore, this work identified opportunities for future exploratory efforts on therapies that might attenuate breast cancer progression by potentially blocking CD44 in circulating tumor bEVs and sEVs or selectively depleting tumor-specific EVs in circulation.

**Figure 9.**
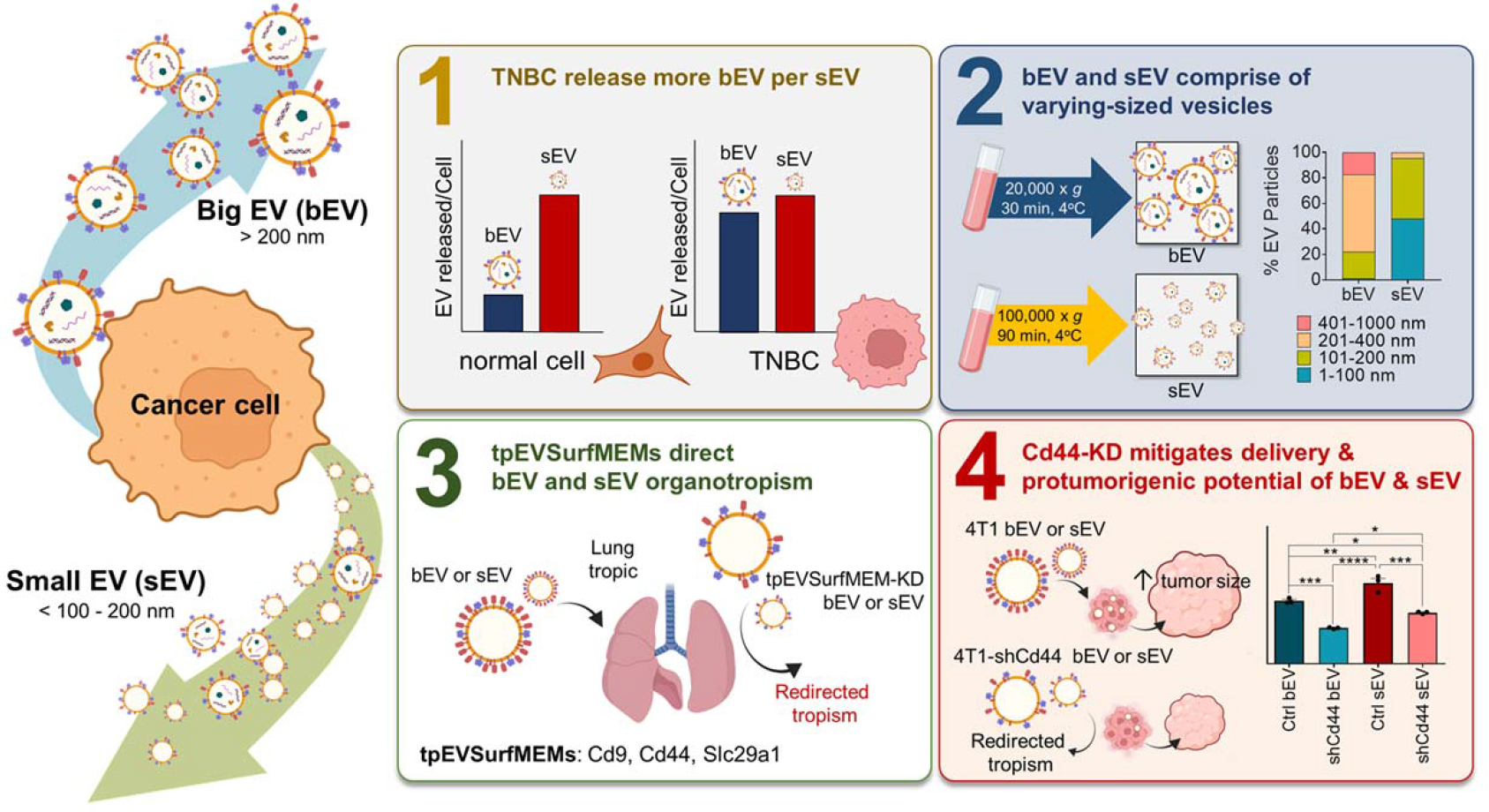
Membrane Protein Modification Modulates Big and Small Extracellular Vesicle Biodistribution and Tumorigenic Potential in Breast Cancers *in vivo*. Graphical summary showing that aggressive TNBC released more bEVs per sEVs than normal mammary epithelial cells. Systematic characterization reveals that TNBC-derived bEVs and sEVs exhibited distinctly different particle size compositions and innate *in vivo* biodistribution profiles. Despite these differences, both bEVs and sEVs demonstrated similar degrees of tropism to the tumor and induced protumorigenic effects upon long-term *in vivo* treatment. Both the tropism to tumor and protumorigenic potential of the bEVs and sEVs were remarkably attenuated upon genetic modification of the EV surface membrane (i.e., depletion of CD44 expression).

## 5. Methods

### 5.1. Cell culture

Murine 4T1 breast cancer cells (gifted by Dr. Yunching Chen, National Tsing Hua University, Hsinchu, Taiwan), human mammary epithelial M10 cells, human TNBC MDA-MB-231, BT-549 (gifted by Dr. Ruey-Hwa Chen, Academia Sinica, Taipei, Taiwan), and Hs578T (Bioresource Collection and Research Center, Hsinchu, Taiwan) cells were cultured in high-glucose Dulbecco’s modified Eagle’s medium (DMEM; HyClone, South Logan, UT, USA) supplemented with fetal bovine serum (FBS; 10% v/v; HyClone), penicillin (100 U mL^-1^; HyClone), and streptomycin (100 µg mL^−1^; HyClone). ER/PR+ (SK-BR-3) and HER2+ (MCF-7) breast cancer cells (gifted by Dr. Ruey-Hwa Chen, Academia Sinica) were maintained in high-glucose DMEM supplemented with FBS (10% v/v), penicillin (100 U mL^-1^), streptomycin (100 µg mL^-1^), and GlutaMAX (1 U mL^−^1; Gibco, Grand Island, NY, USA). All culture media were supplemented with Plasmocin (2.5 µg mL^−1^; InvivoGen, San Diego, CA, USA) to mitigate mycoplasma contamination. Puromycin and Plasmocin were omitted for at least two cell passages prior to any subsequent experiment.

### 5.2. Animal studies

Female BALB/c mice were acquired from BioLASCO Taiwan Co., Ltd. (Ilan, Taiwan) for all *in vivo* experiments. The mice were euthanized by thigh intramuscular injections of Zoletil 50 (200 µL, 25 mg mL^−1^; Sigma-Aldrich, St. Louis, MO, USA) and xylazine (20 mg mL^−1^; Sigma-Aldrich), followed by cervical relocation. All animals included in this work received humane care in compliance with the National Academy of Sciences Guide for the Care and Use of Laboratory Animals. The National Tsing Hua University (NTHU) Animal Research Committee approved all animal-related procedures and protocols (IACUC Protocol No. 108021) performed in this study.

### 5.3. Generation of 4T1-PalmGRET, -shCd9, -shCd44, and -shSlc29a1 stable cell lines and EVs

To generate the 4T1-PalmGRET stable cell line, pLenti CMV Puro DEST plasmids encoding PalmGRET (Plasmid #158221; Addgene, Watertown, MA, USA) were packaged into lentiviruses, then transduced into the 4T1 cells. Non-transduced cells were eliminated by puromycin selection (1 µg mL^−1^, MDbio, Taipei, Taiwan). Additional selection using fluorescence-activated cell sorting was performed to screen for cells that highly expressed PalmGRET.

To generate the 4T1-PalmGRET tpEVSurfMEM KD stable cell lines, pLKO.1 plasmids encoding shRNA targeting mouse *Cd9* (TRCN0000066396), *Cd44* (TRCN0000262945), or *Slc29a1* (TRCN0000079726) or Scramble (ASN0000000002) from the National RNAi Core Facility (Academia Sinica) were packaged into lentiviruses, which were either used directly for experiments or stored in aliquots at −80°C until subsequent experiments. The 4T1-PalmGRET cells were transduced with the packaged lentiviruses to produce stable cell lines with depleted *Cd9*, *Cd44*, or *Slc29a1* expression. The 4T1- PalmGRET-shCd9, -shCd44, -shSlc29a1, and -Scramble cell lines subsequently underwent real-time (RT)-qPCR to examine targeted mRNA expression and were cultured for the KD- EV isolation experiments.

### 5.4. RNA isolation and RT-qPCR

The total RNAs were isolated from stable 4T1- PalmGRET-shCd9, -shCd44, -shSlc29a1, and -Scramble cells using a RNeasy Mini Kit (Qiagen, Hilden, Germany) and underwent RT-qPCR to quantify *Cd9*, *Cd44*, *Slc29a1*, and β- actin mRNA expression. Reverse transcription of the total RNAs into complementary DNA (cDNA) was performed using SuperScript™ IV First-Strand Synthesis System (Invitrogen) before qPCR was performed with SYBR Green PCR Master Mix (Invitrogen) using a StepOnePlus Real-Time PCR System (Applied Biosystem: 95°C for 10 min, 40 cycles at 95°C for 15 s, and 62°C for 1 min). The relative gene expressions were obtained using the comparative threshold cycle (ΔΔCT) method. **Table S3** lists the primers used.

### 5.5. Isolation of bEVs and sEVs

The 4T1, 4T1-PalmGRET, MDA-MB-231, Hs578T, and BT-549 cells were grown in high-glucose DMEM supplemented with EV-depleted FBS (5% v/v), penicillin (100 U mL^-1^), and streptomycin (100 µg mL^-1^). The MCF-7 and SK-BR-3 cells were cultured in high-glucose DMEM supplemented with EV-depleted FBS (5% v/v), penicillin (100 U mL^−1^), streptomycin (100 µg mL^-1^), and GlutaMAX (1 U mL^−1^) for 48 h at 37°C in a humidified incubator with CO_2_ (5%). Conditioned medium (CM) collected from 80–90% confluent cell culture dishes were pooled then subjected to differential centrifugation. The CM were first centrifuged at 300 ×*g* and 2000 ×*g* at 4°C for 10 min to separate cells and cell debris, respectively. Next, the supernatants were centrifuged at 20,000 ×*g* at 4 °C for 30 min, and the bEV pellets were suspended in double 0.22-µm-filtered phosphate-buffered saline (PBS). The remaining supernatants were subsequently centrifuged at 100,000 ×*g* at 4°C for 90 min, and the sEV pellets were suspended in double 0.22-µm- filtered PBS. The harvested bEVs and sEVs were topped up to 5 mL with double 0.22-µm- filtered PBS then centrifuged at 20,000 ×*g* (bEVs) or 100,000 ×*g* (sEVs) at 4°C for 60 min to remove non-EV contaminants. The washed bEV and sEV pellets were suspended in double 0.22-µm-filtered PBS. The protein concentrations of the isolated EVs were determined by a Pierce BCA Kit (Thermo Fisher Scientific, Waltham, MA, USA). The bEV and sEV samples were either used directly for experiments or stored in aliquots at −80°C.

### 5.6. Transmission electron microscopy (TEM)

Isolated bEVs and sEVs (5 µL) were applied onto glow-discharged 400 mesh formvar/carbon-coated grids (Ted, Pella, Redding, CA, USA), then negatively stained with 1% uranyl formate. Dried sample grids were viewed using an FEI Tecnai G2 F20 S-TWIN TEM (Thermo Fisher Scientific) operated at 120 kV. Images were obtained via a Gatan CCD camera and processed using Gatan Digital Micrograph software (Gatan, Inc., Pleasanton, CA, USA). The mean bEV and sEV particle sizes were determined from the analyzed bEV and sEV particles (*N* = 400) detected in the TEM images using Fiji (ImageJ, NIH).

### 5.7. Lipophilic fluorescent dye labelling

Freshly isolated EVs were labeled with PKH26 (Sigma-Aldrich), DiD, or DiR (Thermo Fisher Scientific). PKH26 labeling was performed using the manufacturer’s recommended protocol while DiD and DiR labelling was performed using the manufacturer’s recommended protocol with minor modifications. Briefly, freshly isolated bEV or sEV samples were incubated with DiD or DiR (5 µM) for 20 min at 37°C protected from light. Subsequently, the EV samples were topped up to 5 mL with double 0.22-µm-filtered PBS and centrifuged at 20,000 ×*g* (bEVs) or 100,000 ×*g* (sEVs) at 4°C for 60 min to remove free dye. The washed dyed bEV and sEV pellets were suspended in double 0.22-µm-filtered PBS and were either used directly for experiments or stored in aliquots at −80°C.

### 5.8. NTA

The EV particle sizes and concentrations were determined using a NanoSight NS300 Instrument version 3.2 Dev Build 3.2.16 (Malvern Panalytical, Malvern, UK). Triplicate measurements per sample were made using capture camera level 10 with temperature control at 25 ± 0.01°C.

### 5.9. Sucrose density gradient analysis

Sucrose gradient fractionation was performed by first layering sucrose (8%, 30%, 45%, and 60% w/v; Sigma-Aldrich) in PBS (pH 7.4), then loading the isolated bEVs or sEVs on top of the discontinuous sucrose gradient. Subsequently, the sample was centrifuged at 268,000 ×*g* at 4°C for 38 min with an MLA-50 rotor (Beckman Coulter, Brea, CA, USA). The top layer and subsequent 10 fractions (fractions 1–10) were collected. Aliquots (16.5 µL) of each fraction was saved for BLI assay. Fractions 2–9 were diluted 1:10 in PBS and centrifuged at 20,000 ×*g* (bEVs) or 100,000 ×*g* (sEVs) at 4°C for 1 h with an MLS-55 rotor (Beckman Coulter). The bEV and sEV pellets were suspended in double 0.22-µm-filtered PBS and were either used directly for experiments or stored in aliquots at −80°C until subsequent BLI, NTA, or western blotting analyses.

### 5.10. Western blotting

The cells and EVs were lysed in radioimmunoprecipitation assay lysis buffer (1× PBS, pH 7.4, with 1% v/v Igepal CA-630, 0.5% w/v sodium deoxycholate, and 0.1% w/v sodium dodecyl sulfate) with cOmplete protease inhibitor cocktail (Roche, Basel, Switzerland). The EV and cell lysate protein concentrations were determined using a Pierce BCA Protein Assay Kit (Thermo Fisher Scientific). The protein lysates were separated using Bolt 4–12% Bis-Tris Plus Gels (Burlington, MA, USA) and transferred onto 0.45-µm polyvinylidene fluoride membranes (Amersham Bioscience, Little Chalfont, UK). The membranes were blocked with bovine serum albumin (BSA; 5%; Cyrusbioscience, Taipei, Taiwan) in Tris-buffered saline (TBS; pH 7.4) with Tween 20 (0.1% v/v; TBST) for 1 h at room temperature. The blots were immunoprobed with primary antibodies overnight at 4°C, washed thrice in TBST at room temperature, then incubated with infrared (IR) dye-conjugated secondary antibodies for 1 h at room temperature. The blots were imaged using an Odyssey CLx unit (LI-COR Biosciences, Lincoln, NE, USA). **Table S6** lists the information on the primary and secondary antibodies used.

### 5.11. Dot blot analysis

The purified EVs were serially diluted to 2000, 1000, 500, and 250 ng in 5 µL 0.22-µm-filtered PBS to characterize the PalmGRET membrane orientation in the bEVs and sEVs. The cell lysates were diluted to 1000 ng in 5 µL 0.22-µm-filtered PBS as positive controls. The diluted bEVs, sEVs, cell lysates, and double 0.22-µm-filtered PBS (all, 5 µL) were dotted onto 0.22-µm nitrocellulose membranes (Amersham Bioscience) and blocked in either TBS (5% BSA w/v) or TBST (5% BSA w/v) overnight at 4°C. The membranes were immunoprobed with anti-GFP antibody (GeneTex, Irvine, CA, USA) diluted in either TBS (5% BSA w/v) or TBST (5% BSA w/v) overnight at 4°C, washed thrice in TBS or TBST at 30 min per wash, incubated with IR dye-conjugated secondary antibodies for 1 h at room temperature, and imaged by an Odyssey CLx unit (LI-COR Biosciences). **Table S6** lists the information on the primary and secondary antibodies used.

### 5.12. *In vivo* and *ex vivo* imaging

BALB/c mice were anesthetized after bolus injection of 4T1-PalmGRET-DiR bEV or sEV (100 µg). Subsequently, diluted Fz (1/40; Promega, Madison, WI, USA) in 200 µL 0.22 µm-filtered PBS was IV-injected to the mice via the tail vein for IVIS imaging. Images were captured at 15 min, 4 h, and 48 h post-Fz administration. Following initial brightfield imaging, PalmGRET signals (via BRET-FL) were obtained using a 520/20 nm emission filter. A 740/20-nm excitation filter and 800/20-nm emission filter were used to acquire DiR signals. Following live animal imaging, each mouse was anesthetized again and euthanized with cervical dislocation. Major organs (the brain, lungs, heart, liver, spleen, kidneys, stomach, intestine, spine, femur, and fat pad) were collected for *ex vivo* imaging. The harvested organs were immersed in diluted Fz (1/400; Promega) in sterile PBS then imaged using IVIS (Perkin Elmer, Massachusetts, USA) to detect PalmGRET (via BRET-FL; 520 nm emission filter) and DiR (via FL excitation; 740 nm excitation/800 nm emission filters) signals.

### 5.13. MS and proteomics data analysis

Polyacrylamide gel slices containing bEV and sEV protein lysates were resolved in tris (2-carboxyethyl) phosphine (10 × 10^-1^ M; Sigma-Aldrich) with ammonium bicarbonate (50 × 10^-1^ M; Sigma-Aldrich) for 30 min at 37°C and alkylated in iodoacetamide (50 × 10^-1^ M; Sigma-Aldrich) with ammonium bicarbonate (50 × 10^-1^ M) for 45 min in the dark at 25°C. Trypsin/Lys-C solution (Promega) was added and incubated at 37°C for 12 h. The resulting peptides were extracted from the gel fragments and analyzed with an Orbitrap Fusion Lumos Tribrid quadrupole-ion trap-Orbitrap mass spectrometer (Thermo Fisher Scientific) combined with an UltiMate 3000 nanoLC system (Thermo Fisher Scientific, Bremen, Germany) with higher energy collision dissociation (HCD) MS/MS mode.

MaxQuant version 1.6.5.01[32] was used to obtain the secondary MS data utilizing the Swiss-Prot *Mus musculus* database (Swiss-Prot, 17,009 reviewed protein sequences, UP000000589) for peptide identification. The criteria for the database searches were as follows: carboxyamidomethylation on cysteine (fixed modification), oxidation of methionine (variable modification), minimum of five amino acids in the peptide length, precursor mass tolerance of 20 ppm, and fragment ion tolerance of 0.5 Da. A false discovery rate of 0.01 and peptide rank of 1 were set for the peptide identification filters. The MS proteomics data have been deposited into the ProteomeXchange Consortium via the PRIDE[33] partner.

### 5.14. Bioluminescence assays

Nluc activity was measured using a GloMax Discover Microplate Reader (Promega) coupled with auto-injectors and set for Nluc (450/10 nm) and BRET-FL (510/10 nm) readings. The BRET ratio was computed as the quotient of the acceptor signal (510/10 nm) by the donor signal (450/10 nm). To determine EV bioluminescence (BL) activity, triplicate samples of EVs (5 µL), crude sucrose fractions (5 µL), or pelleted sucrose fractions (3 µL) were loaded onto a 96-well white plate (Greiner Bio-One, Monroe, NC, USA). The Nluc and BRET signals were measured (integration time: 1 s) after auto-injecting either Fz (Promega) or FFz (Promega) in PBS (25 µM final concentration). For EV biodistribution analysis, the collected organs were homogenized and lysed using T-PER (0.5 g organ mL^−1^ T-PER, Thermo Fisher Scientific) supplemented with 1× Halt protease inhibitor (Thermo Fisher Scientific). The lysates were centrifuged at 10,000 ×*g* at 4°C for 10 min, followed by supernatant collection to acquire the protein extracts. The supernatants (20 µL per well) were loaded into a 96-well white plate (Greiner Bio-One) in triplicate. The BL signals were measured after auto-injection of FFz (80 µL) in PBS per well (25 µM final concentration) with a 1-s integration time.

### 5.15. EV organotropism redirection study by EV membrane modification

BALB/c mice were injected IV with 4T1-PalmGRET-shCd9-, -shCd44-, -shSlc29a1-, or -Scramble- bEVs or -sEVs (10 µg) at 0, 24, 48, and 72 h through the tail vein. At 0.5 h and 72.5 h post-injection, the mice were anesthetized, euthanized by cervical dislocation, and dissected to obtain the blood (via cardiac puncture) and organs (brain, lungs, heart, liver, spleen, kidney, stomach, spinal cord, femur, fat pad) for BLI biodistribution analyses. The total RLU per organ was determined as follows: (RLU/20 µL) × (organ mL^−1^ T-PER lysis buffer). The proportions of the detected EV signal were computed as follows: [(RLU of specific organ)/(RLU of all organs)] × 100%.

### 5.16. Murine breast cancer model

Murine 4T1 breast cancer cells were cultured, harvested, and suspended in PBS (pH 7.4). BALB/c mice were anesthetized prior to cancer cell injection. The breast cancer tumor model was established by subcutaneous injection of 4T1 cells (5 × 10^4^ cells in 100 µL PBS) into the mammary fat pads of 6–8-week-old female BALB/c mice.

### 5.17. Immunohistochemistry

For histological analysis, the tissues were dissected, fixed in paraformaldehyde (4% w/v), and embedded in paraffin. Subsequently, the tissue blocks were cut into sections (5-μm). Heat-mediated antigen retrieval with citrate buffer (pH 6.0) was performed on the deparaffinized and rehydrated tissue sections. Next, the sections were washed with TBS-Triton® X-100 (0.025% v/v), then blocked with TBS containing BSA (1% w/v) and normal goat serum (2% v/v) for 1 h at room temperature. The sections were incubated with primary antibodies in a humidified chamber at 4°C overnight, washed with TBS-Triton® X-100 (0.025% v/v), and incubated with Alexa Fluor 568-conjugated secondary antibody for 1 h at room temperature. Subsequently, autofluorescence was quenched using a Vector® TrueVIEW® Autofluorescence Quenching Kit (Vector Laboratories, Newark, CA, USA). The sections were washed with PBS and mounted using VECTASHIELD® Vibrance™ mounting medium with 4′,6-diamidino-2-phenylindole (DAPI, Vector Laboratories). **Table S6** lists the primary and secondary antibody dilutions for the organ tissues. All tissue sections were imaged using an LSM 980 Airyscan 2 confocal laser scanning microscope (Zeiss, Stuttgart, Germany) under similar imaging parameters. The images were acquired and processed using ZEN blue 3.3 (Zeiss). EV quantitative analyses were performed using Fiji (ImageJ) under similar threshold settings.

### 5.18. Statistical analyses

Statistical analyses were performed using GraphPad Prism version 7.05. A 2-tailed Student’s *t*-test was used for comparison between two groups. A one-way analysis of variance (ANOVA) followed by multiple group comparisons was used for comparison of ≥3 groups. ANOVA followed by Dunnett’s post hoc test was used to compare experimental groups with the control. The values were normally distributed and the variance was similar between compared groups. Error bars for all graphs represent the mean ± SEM. *P* < 0.05 was considered statistically significant (n.s., not significant).

## Supporting Information

Supporting Information is available from the Wiley Online Library or from the author.

## Acknowledgements

The authors thank Dr. Wei Chun Huang (Biophysics Core Facility, Institute of Atomic and Molecular Sciences, Academia Sinica) for assistance and technical support, Chia Chen Tai and Tzu Wen Tai [Flow Cytometry Core Facility, Academia Sinica Core Facility and Innovative Instrument Project (AS-CFII 108-113)] for assistance on the cell sorting service, the National RNAi Core Facility (Academia Sinica) for providing the shRNA plasmids, Dr. Hsin-Yi Wu (NTU Consortia of Key Technologies and NTU Instrumentation Center) for providing mass spectrometry technical research services, and the NTHU Biomedical Science & Engineering Center Animal Care Facility; and Academia Sinica Biological Electron Microscopy Core Facility, Academia Sinica Core Facility and Innovative Instrument Project (AS-CFII-111-203). The Lai Lab was supported by the National Science and Technology Council (NSTC) under Grant Nos. 111-2628-B-001-004 (C.P.L.) and 111-2628-B-001-022 (C.P.L.), Academia Sinica Innovative Materials and Analysis Technology Exploration (iMATE) Program AS-iMATE-111-34 (C.P.L.), Academia Sinica Career Development Award AS-CDA-109-M04 (C.P.L.). Anthony Yan-Tang Wu and Yen-Ju Chen contributed equally to this work.

Received: ((will be filled in by the editorial staff))

Revised: ((will be filled in by the editorial staff))

Published online: ((will be filled in by the editorial staff))

## Table of Contents

Bryan John Abel Magoling, Anthony Yan-Tang Wu, Yen-Ju Chen, Wendy Wan-Ting Wong, Steven Ting-Yu Chuo, Hsi-Chien Huang, Yun-Chieh Sung, Hsin Tzu Hsieh, Poya Huang, Kang-Zhang Lee, Kuan-Wei Huang, Ruey-Hwa Chen, Yunching Chen, Charles Pin-Kuang Lai*

Membrane Protein Modification Modulates Big and Small Extracellular Vesicle Biodistribution and Tumorigenic Potential in Breast Cancers *in vivo*

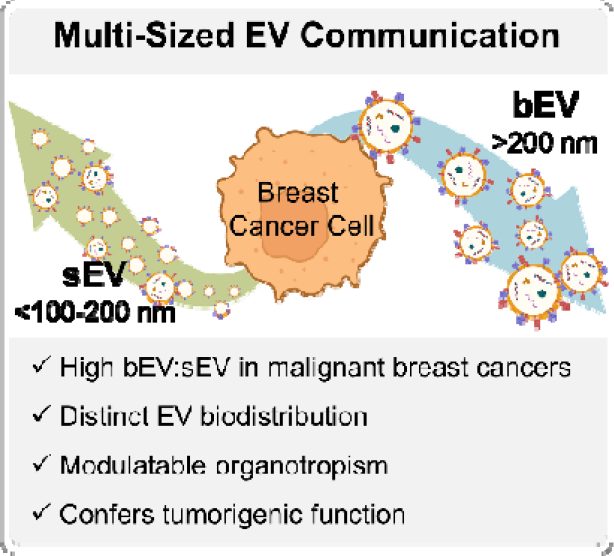

**Big and small extracellular vesicles circulate differentially and confer tumorigenic function modulatable by EV membrane proteins.** Malignant breast cancers release more bEV, in addition to sEVs, than normal cells to target organs and individually promote tumor growth *via* tumor progression-associated EV surface membrane proteins (tpEVSurfMEMs).

## Supporting Information

**Supporting Figure 1.**
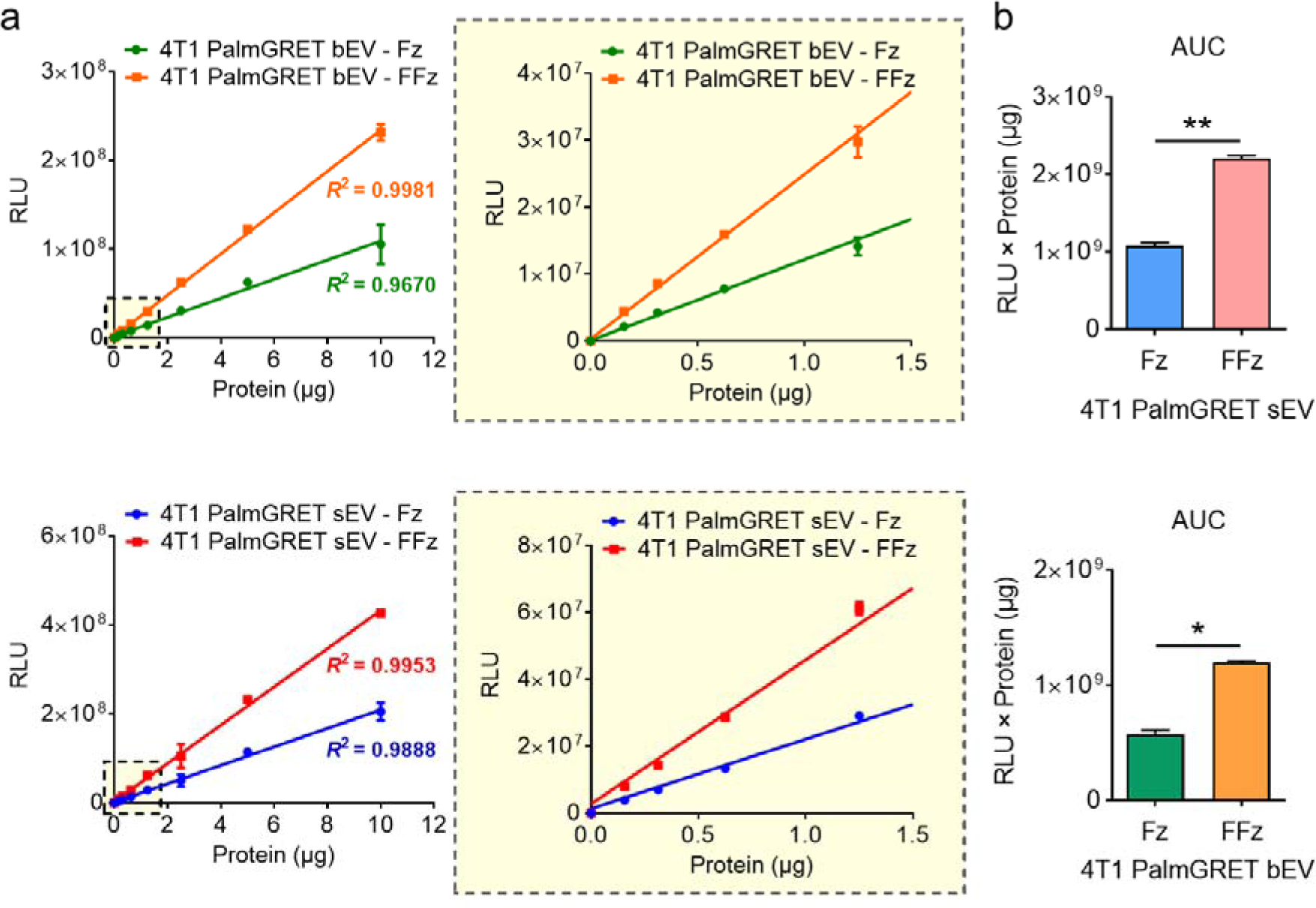
FFz yields a higher BLI signal in PalmGRET EVs. a, The plot of Nluc activity vs. protein of bEVs and sEVs showing significantly increased BLI signals when FFz was applied as compared to Fz. Both Fz- and FFz-induced Nluc signals were detected in an EV protein amount-dependent manner and with high correlation in both bEV and sEV samples (*R^2^* ≥∼0.99)**. b,** Area under the curve (AUC) bar graphs of (**a**) demonstrate that FFz yielded Nluc signals that were ∼2-fold higher than that induced by Fz. \**p* < 0.05; \*\**p* < 0.01 with 2-tailed Student’s *t*-test.

**Supporting Figure 2.**
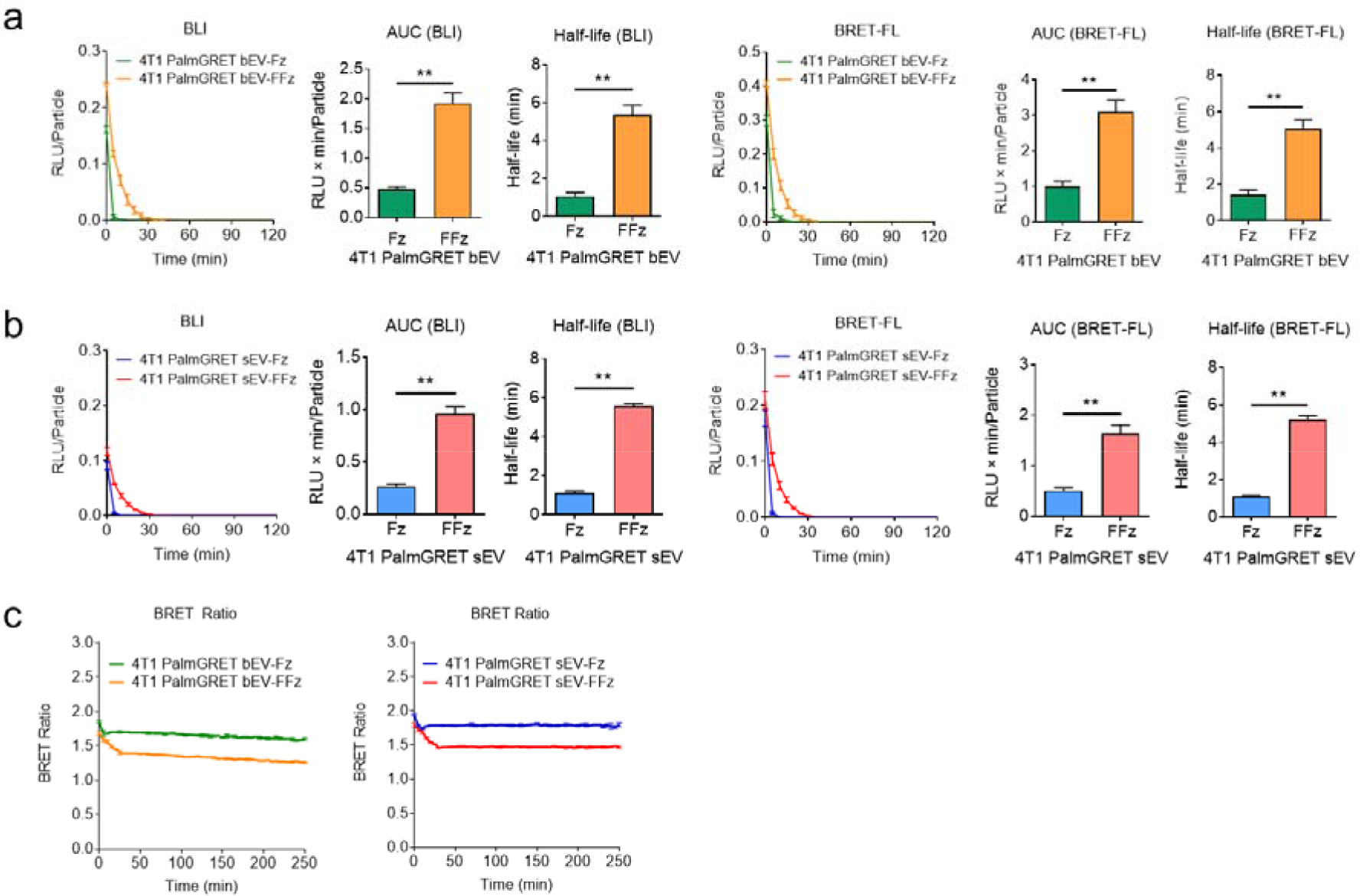
FFz yields a longer half-life of BLI and BRET-FL signals in PalmGRET EVs. The BLI and BRET-FL of **a,** bEVs and **b,** sEVs demonstrated significantly increased BLI half-life when FFz was used as compared to Fz. **c,** The BRET ratio of bEVs and sEVs when reacted with FFz and Fz. Compared to Fz-induced activity, the BRET ratio of FFz-induced signals decreased steadily for the first 30 min before plateauing, which suggested increased enzyme–substrate interaction between Nluc and FFz and hence greater substrate consumption over time than Fz. \*\**p* < 0.01 with 2-tailed Student’s *t*-test.

**Supporting Figure 3.**
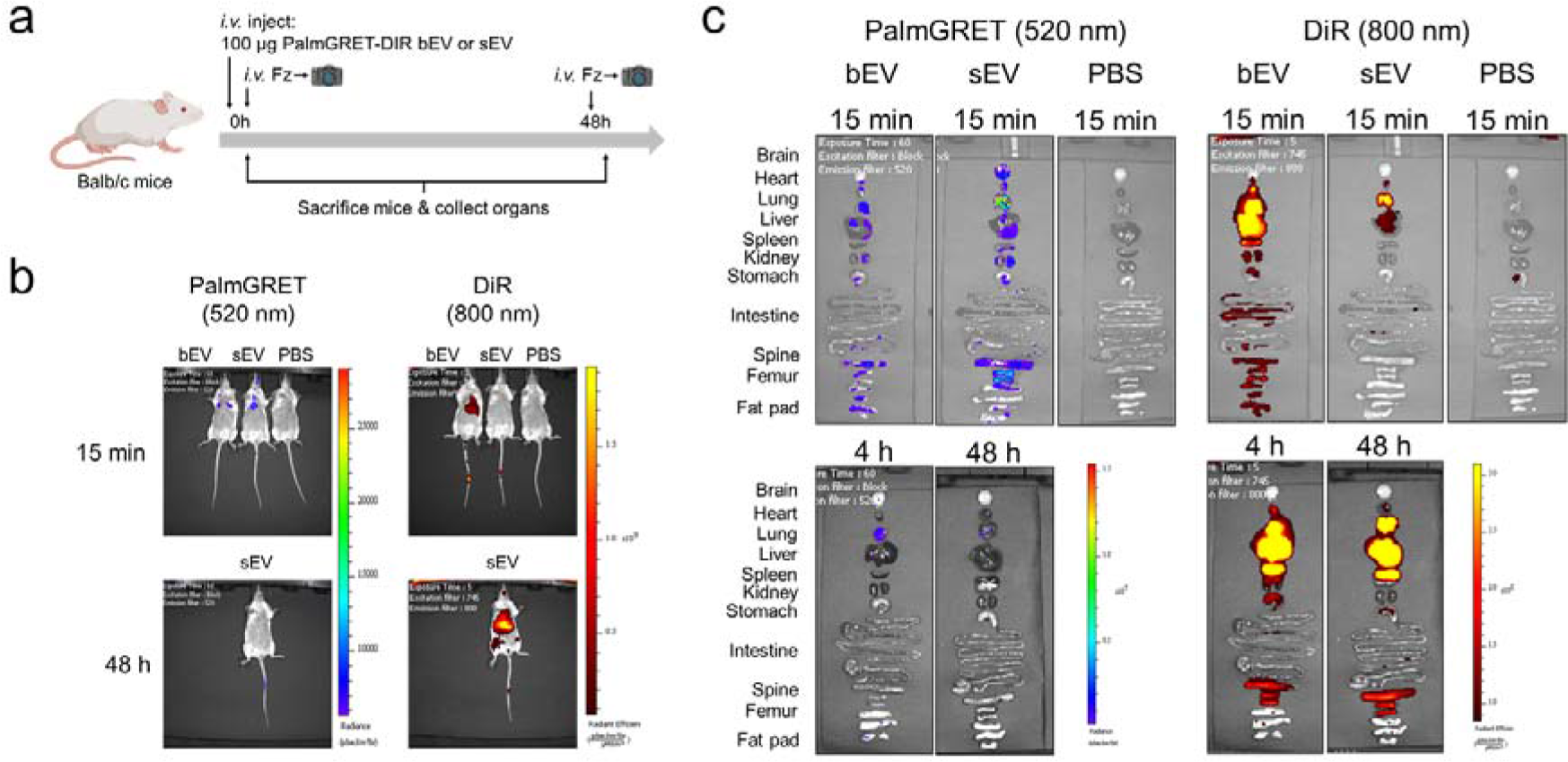
*In vivo* imaging of 4T1-PalmGRET-DiR-bEVs and -sEVs. a, Schematic of live animal and *ex vivo* imaging of BALB/c mice injected IV with 4T1- PalmGRET-DiR-bEVs and -sEVs. **b,** *In vivo* and **(c)** *ex vivo* images of BALB/c mice administered with 4T1-PalmGRET-DiR-bEVs and -sEVs at 15 min, 4 h, and 48 h post-injection.

**Supporting Figure 4.**
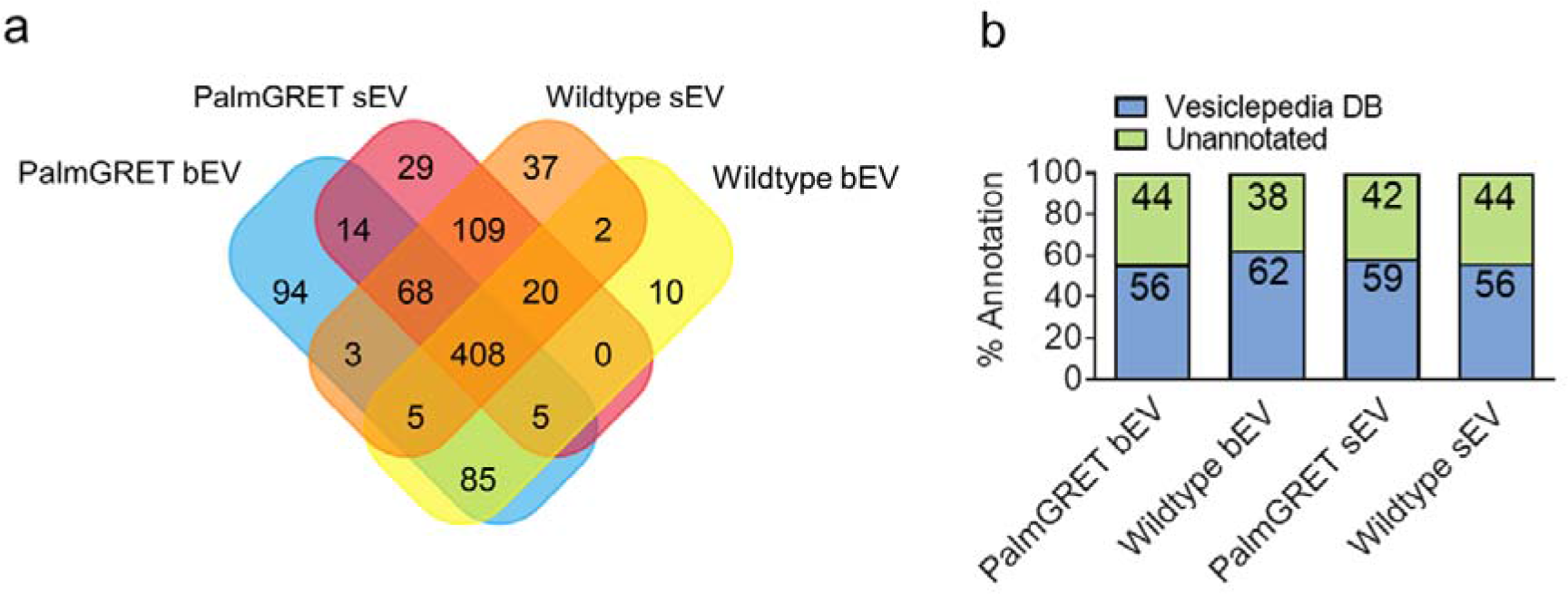
Proteomic analysis of bEVs and sEVs derived from TNBC 4T1 cells. a, Venn diagram showing unique and common proteins between 4T1-PalmGRET-bEVs and -sEVs and 4T1-WT-bEV and -sEV proteomes. **b,** Bar graph showing the percentage of proteins in each proteome group previously reported in the Vesiclepedia EV database[34].

**Supporting Figure 5.**
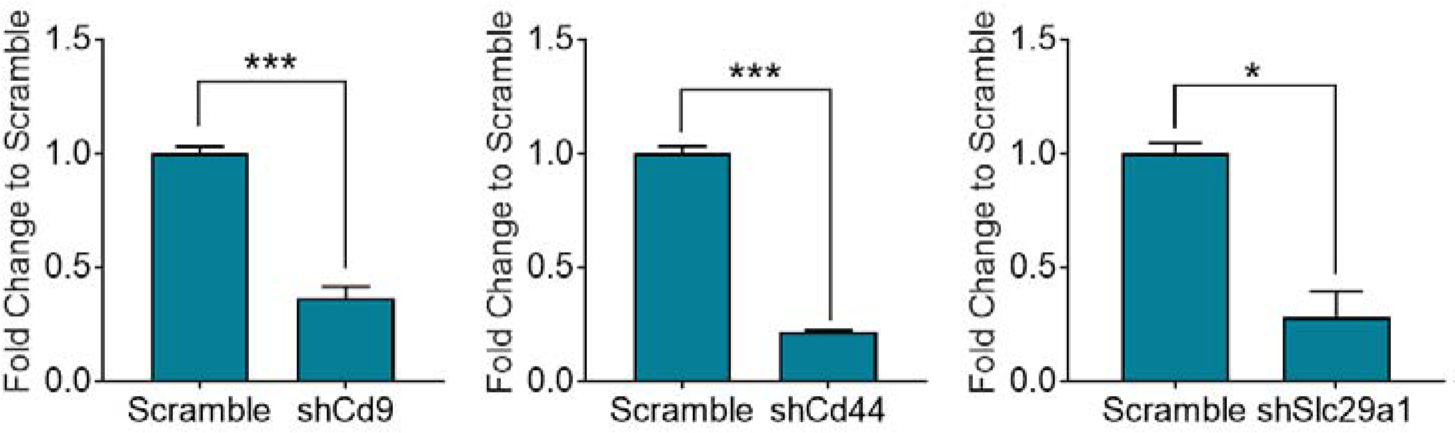
tpEVSurfMEM transcript KD in 4T1-PalmGRET cells. RT-qPCR indicating reduced relative cDNA expression of *CD9*, *CD44*, and *SLC29A1* in the shCd9, shCd44, and shSlc29a1 groups, respectively, as compared to the Scramble controls. \**p* < 0.05; \*\*\**p* < 0.001 with the 2-tailed Student’s *t*-test.

**Supporting Figure 6.**
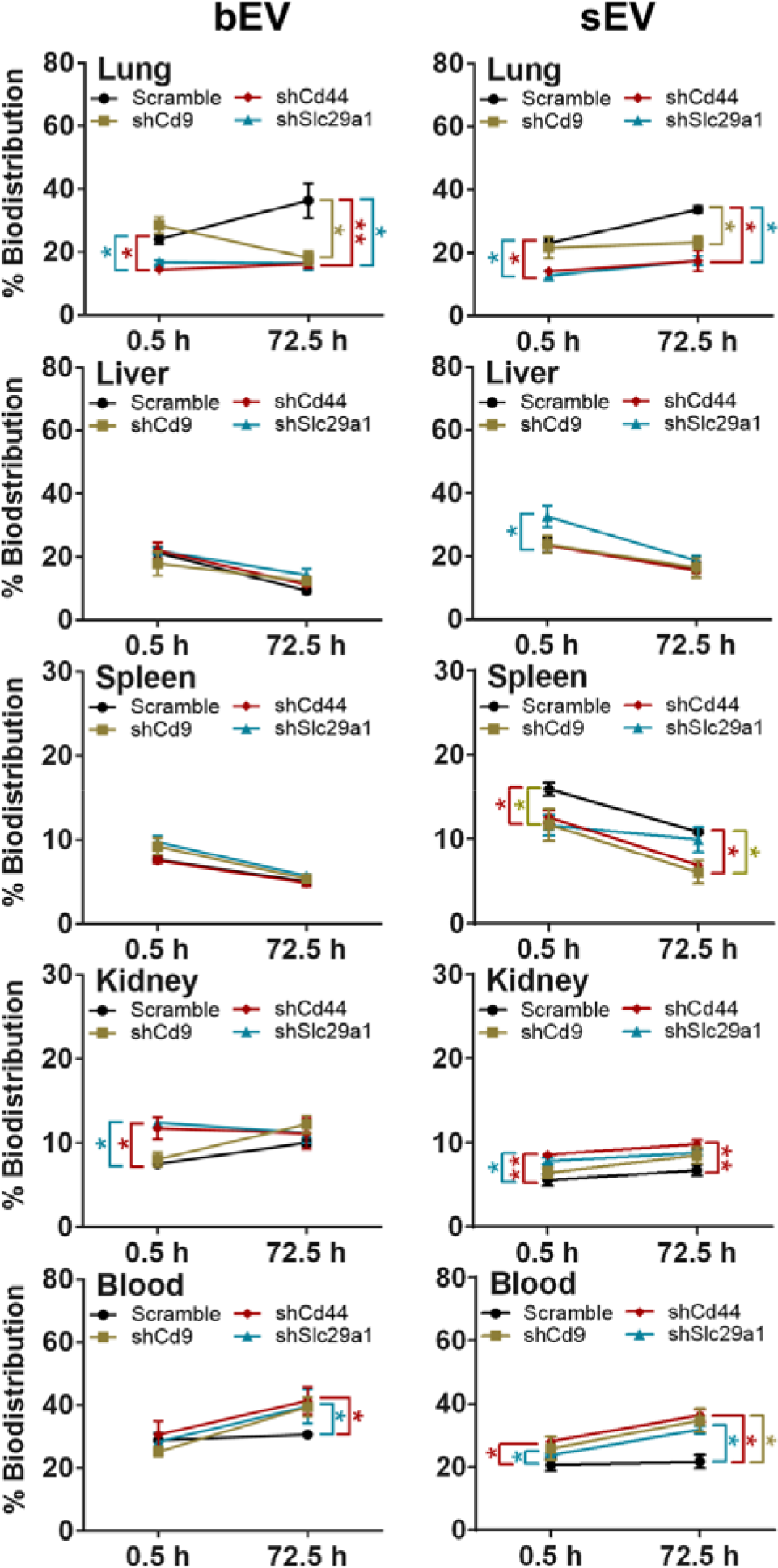
Biodistribution percentage of 4T1-PalmGRET with tpEVSurfMEM KDs in non-tumor bearing mice. Biodistribution percentage of IV-injected shCd9-, shCd44-, and shSlc29a1-4T1-PalmGRET-bEV (left) and -sEV (right) distribution in the lung, liver, kidney, spleen, and blood as compared to the Scramble controls. \**p* < 0.05; \*\**p* < 0.01 with 1-way ANOVA followed by Dunnett’s post hoc test vs. Scramble controls.

**Supporting Figure 7.**
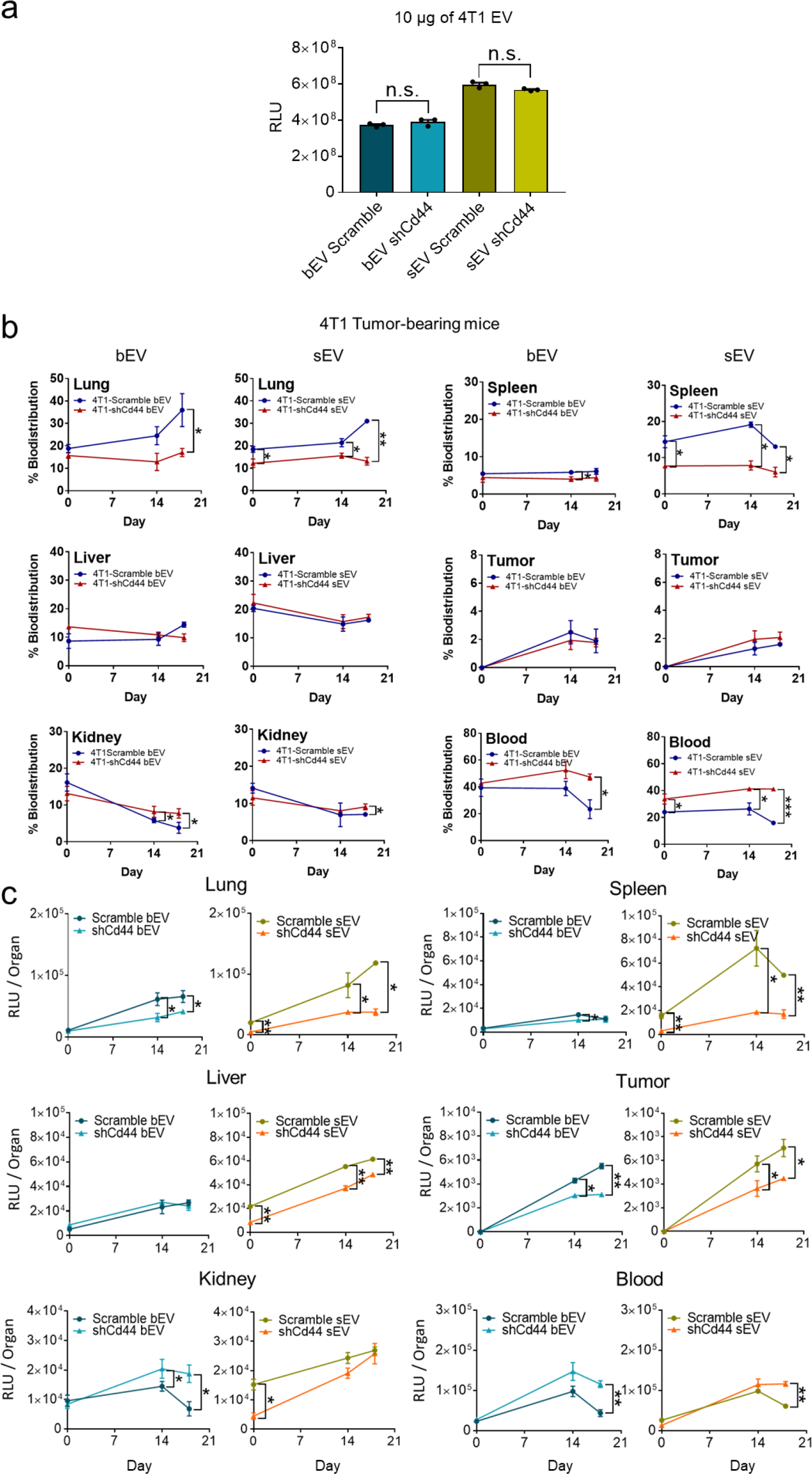
BLI intensity and biodistribution percentage of 4T1-PalmGRET- EVs with Scramble or shCd44 in 4T1-tumor bearing mice. a, Initial BLI intensity was similar between IV-administered shCd44 and Scramble 4T1-PalmGRET-bEVs and sEVs. **b,** Biodistribution percentage of IV-injected 4T1-PalmGRET-shCd44-bEVs (left) and -sEVs (right) as compared with Scramble controls. \**p* < 0.05; \*\**p* < 0.01 with 2-tailed Student’s *t*- test. c) Total EV signal per organ of IV-injected 4T1-PalmGRET-shCd44-bEVs and -sEVs as compared with the Scramble controls. \**p* < 0.05; \*\**p* < 0.01; \*\*\**p* < 0.001 with 2-tailed Student’s *t*-test.

**Supporting Figure 8.**
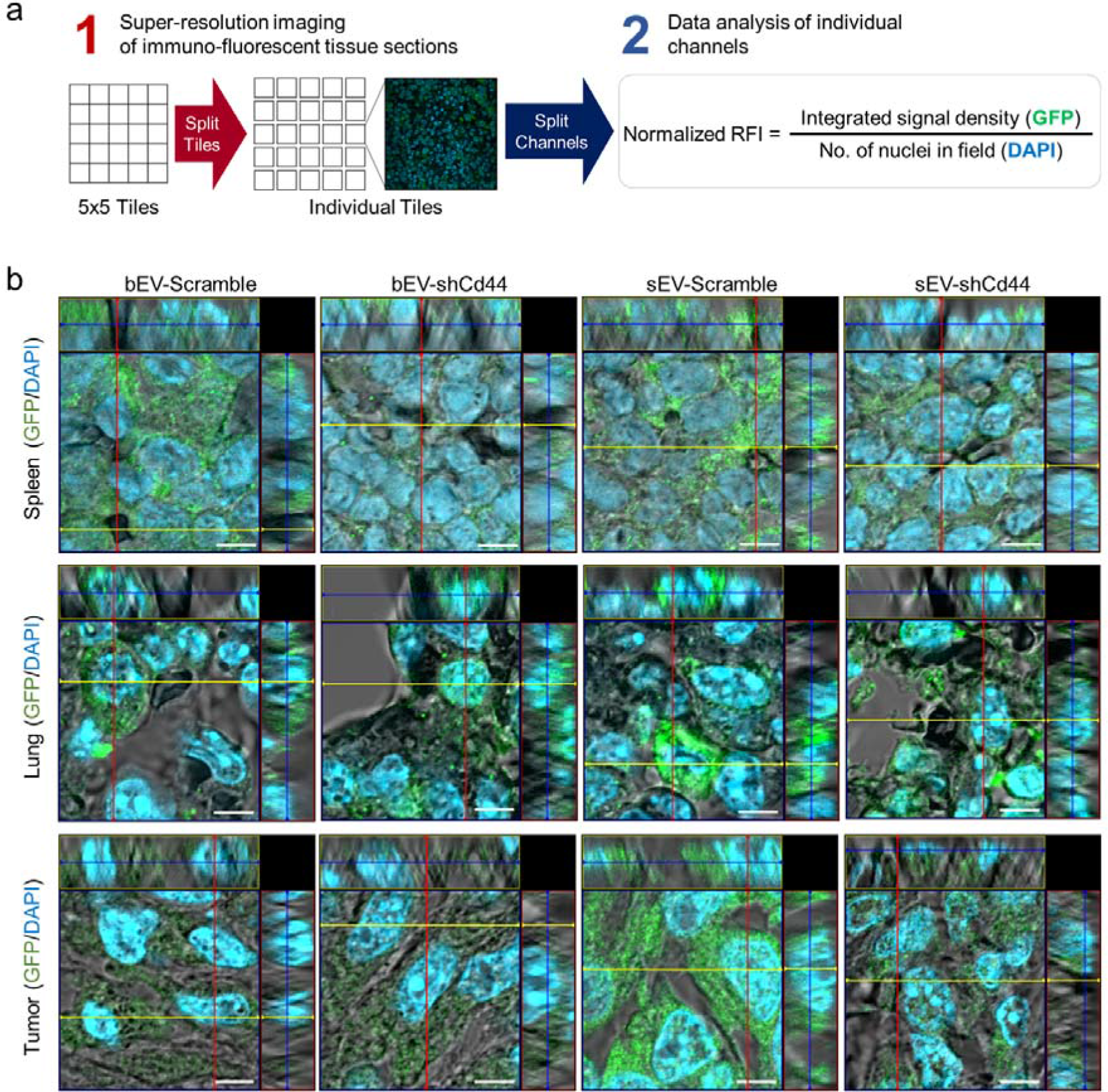
Cellular uptake of 4T1-PalmGRET-EVs shCd44 with Scramble or shCd44 in 4T1-tumor bearing mice. **a**, schematic for image processing of super-resolution images of immuno-fluorescent tissue sections. **b,** Orthogonal view of Z-stack images showing intracellular uptake of bEVs and sEVs in the spleen, lung, and tumor of mice treated with 4T1-PalmGRET-Scramble or -shCd44-bEVs or -sEVs at day 18 post-EV treatment. The nuclei were stained by DAPI and 4T1-PalmGRET was immunoprobed by anti-GFP antibody followed by Alexa Fluor 568-conjugated secondary antibody. Bar, 5 µm.

**Supporting Table 1.**
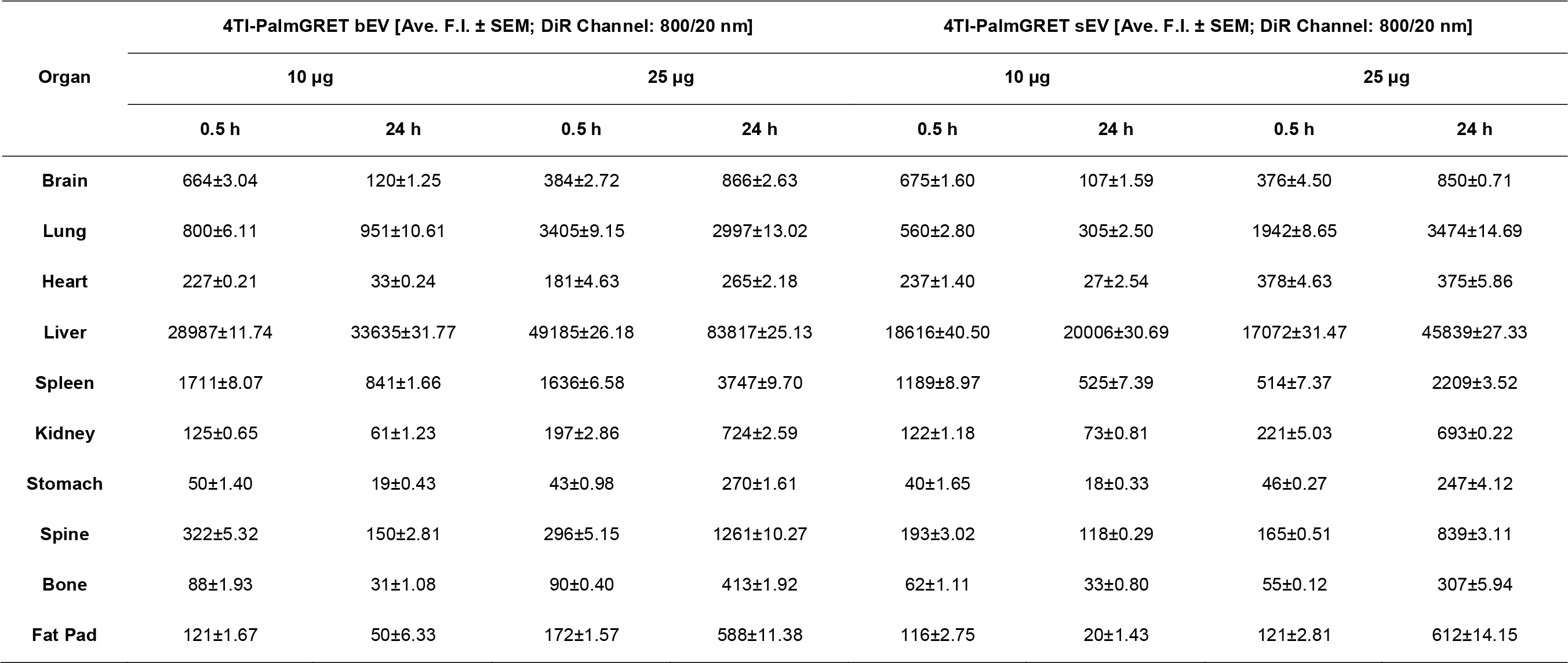
Fluorescence intensity (DiR Channel: 800/20 nm) of 4T1-PamGRET-DiR bEV and sEV at 0.5 h and 24 h post-injection.

**Supporting Table 2.** Organ biodistribution (%) of 4T1-PamGRET-DiR bEV and sEV at 0.5 h and 24 h post-injection.

| Organ | Time | 4T1-PalmGRET-DiR bEV [%] |  |  |  | 4T1-PalmGRET-DiR sEV [%] |  |  |  |
| --- | --- | --- | --- | --- | --- | --- | --- | --- | --- |
|  |  | 10 µg |  | 25 µg |  | 10 µg |  | 25 µg |  |
|  |  | PalmGRET (520 nm) | DiR (800 nm) | PalmGRET (520 nm) | DiR (800 nm) | PalmGRET (520 nm) | DiR (800 nm) | PalmGRET (520 nm) | DiR (800 nm) |
| Brain | 0.5 h | 4.84 | 2.01 | 1.02 | 0.69 | 4.15 | 3.10 | 0.2 | 1.80 |
|  | 24 h | 0.34 | 0.33 | 0.1 | 0.91 | 0.06 | 0.50 | 0.12 | 1.53 |
| Lung | 0.5 h | 32.8 | 2.42 | 51.3 | 6.13 | 36.63 | 2.57 | 53.9 | 9.30 |
|  | 24 h | 7.13 | 2.65 | 4.21 | 3.16 | 1.15 | 1.43 | 8.64 | 6.27 |
| Heart | 0.5 h | 0.99 | 0.68 | 6.96 | 0.32 | 4.37 | 1.08 | 14.78 | 1.81 |
|  | 24 h | 0.26 | 0.09 | 0.18 | 0.28 | 0.07 | 0.13 | 0.95 | 0.68 |
| Liver | 0.5 h | 32.8 | 87.59 | 30.39 | 88.48 | 28.39 | 85.36 | 27.94 | 81.72 |
|  | 24 h | 69.24 | 93.72 | 85.23 | 88.28 | 89.16 | 94.22 | 80.73 | 82.68 |
| Spleen | 0.5 h | 7.02 | 5.17 | 2.49 | 2.94 | 21.47 | 5.45 | 1.24 | 2.46 |
|  | 24 h | 4.45 | 2.34 | 1.97 | 3.95 | 4.21 | 2.47 | 2.7 | 3.98 |
| Kidney | 0.5 h | 3.28 | 0.38 | 1.66 | 0.35 | 3.03 | 0.56 | 0.37 | 1.06 |
|  | 24 h | 10.53 | 0.17 | 5.1 | 0.76 | 1.41 | 0.34 | 3.3 | 1.25 |
| Stomach | 0.5 h | 0.65 | 0.15 | 0.11 | 0.08 | 0.38 | 0.18 | 0.18 | 0.22 |
|  | 24 h | 1.11 | 0.05 | 0.16 | 0.28 | 0.19 | 0.09 | 0.26 | 0.45 |
| Spinal cord | 0.5 h | 11.5 | 0.97 | 4.33 | 0.53 | 9.95 | 0.88 | 1.04 | 0.79 |
|  | 24 h | 2.7 | 0.42 | 1.42 | 1.33 | 2.19 | 0.55 | 1.72 | 1.51 |
| Bone | 0.5 h | 2.65 | 0.27 | 0.68 | 0.16 | 2.9 | 0.29 | 0.14 | 0.27 |
|  | 24 h | 0.94 | 0.09 | 0.76 | 0.43 | 1.3 | 0.16 | 0.87 | 0.55 |
| Fat Pad | 0.5 h | 3.47 | 0.37 | 1.04 | 0.31 | 3.74 | 0.53 | 0.22 | 0.58 |
|  | 24 h | 3.31 | 0.14 | 0.87 | 0.62 | 0.26 | 0.10 | 0.71 | 1.10 |

**Supporting Table 3.**
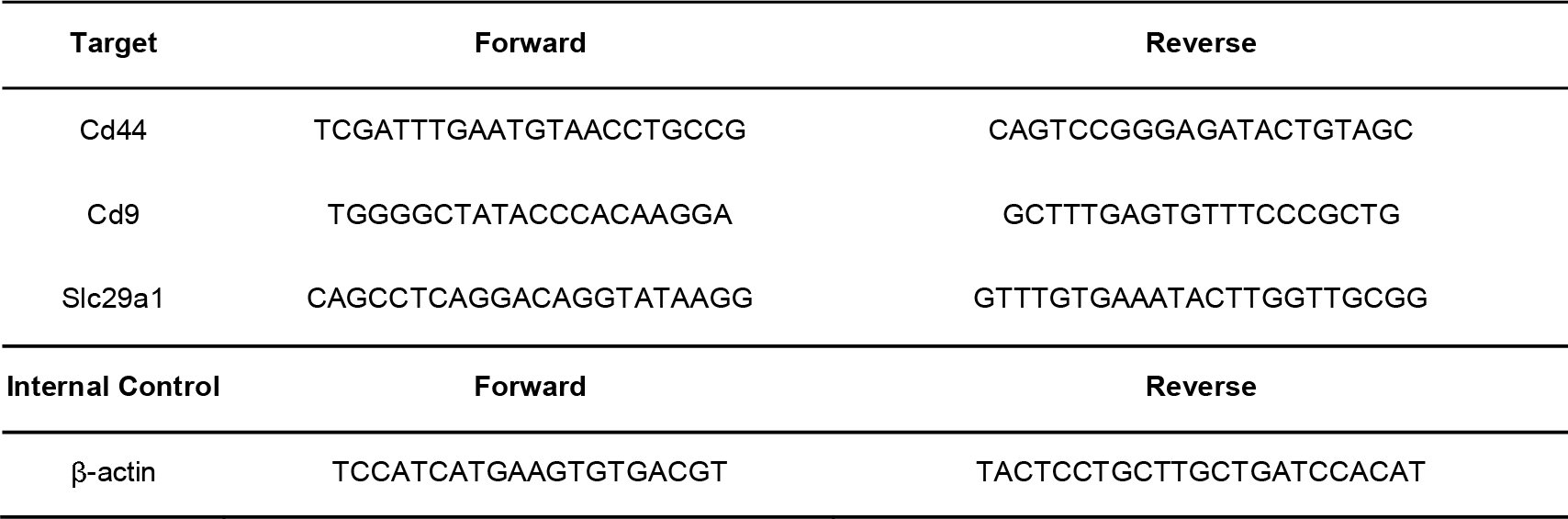
List of primers.

**Supplementary Table 4.** Organ biodistribution (%) of 4T1-PamGRET-Scramble, -shCd9, -shCd44, and -shSlc29a1 bEV and sEV.

| Organ | 10 µg Scramble |  |  |  | 10 µg shCd9 |  |  |  | 10 µg shCd44 |  |  |  | 10 µg shSlc29a1 |  |  |  |
| --- | --- | --- | --- | --- | --- | --- | --- | --- | --- | --- | --- | --- | --- | --- | --- | --- |
|  | bEV [%] |  | sEV [%] |  | bEV [%] |  | sEV [%] |  | bEV [%] |  | sEV [%] |  | bEV [%] |  | sEV [%] |  |
|  | 0.5 h | 72.5 h | 0.5 h | 72.5 h | 0.5 h | 72.5 h | 0.5 h | 72.5 h | 0.5 h | 72.5 h | 0.5 h | 72.5 h | 0.5 h | 72.5 h | 0.5 h | 72.5 h |
| **Brain** | 0.46 | 0.48 | 0.37 | 0.31 | 0.19 | 0.44 | 0.36 | 0.37 | 0.49 | 0.46 | 0.40 | 0.39 | 0.35 | 0.38 | 0.25 | 0.34 |
| **Lung** | 24.06 | 31.71 | 22.98 | 33.74 | 28.46 | 18.16 | 21.63 | 23.25 | 14.49 | 16.27 | 14.18 | 17.41 | 16.80 | 16.43 | 12.79 | 17.55 |
| **Heart** | 0.93 | 1.25 | 0.74 | 1.02 | 0.76 | 1.24 | 0.98 | 0.88 | 1.26 | 1.41 | 1.07 | 1.49 | 0.80 | 1.24 | 0.69 | 1.08 |
| **Liver** | 21.30 | 9.30 | 23.54 | 15.72 | 17.89 | 12.25 | 23.78 | 16.35 | 22.01 | 11.15 | 23.47 | 15.40 | 21.97 | 14.22 | 32.56 | 18.62 |
| **Spleen** | 7.65 | 5.03 | 15.94 | 10.86 | 9.24 | 5.40 | 11.75 | 6.09 | 7.59 | 4.88 | 12.67 | 6.98 | 9.72 | 5.72 | 11.63 | 9.98 |
| **Kidney** | 7.47 | 10.04 | 5.58 | 6.76 | 8.08 | 12.24 | 6.41 | 8.48 | 11.78 | 11.15 | 8.55 | 9.84 | 12.42 | 11.17 | 7.82 | 8.81 |
| **Stomach** | 1.41 | 2.38 | 1.19 | 1.75 | 1.51 | 2.62 | 1.59 | 2.09 | 1.42 | 2.60 | 1.60 | 2.33 | 1.96 | 2.99 | 1.44 | 2.00 |
| **Spinal Cord** | 2.71 | 2.96 | 3.49 | 2.93 | 2.77 | 2.62 | 2.38 | 2.52 | 3.28 | 3.94 | 3.11 | 3.12 | 2.55 | 2.47 | 3.50 | 3.28 |
| **Bone** | 3.47 | 3.90 | 4.35 | 3.73 | 3.93 | 3.44 | 3.91 | 3.40 | 4.77 | 4.12 | 4.87 | 4.25 | 3.58 | 3.25 | 4.03 | 4.47 |
| **Fat Pad** | 1.71 | 2.35 | 1.21 | 1.53 | 1.92 | 2.18 | 1.37 | 1.88 | 2.26 | 2.57 | 2.05 | 2.45 | 1.52 | 2.47 | 1.53 | 2.14 |
| **Blood** | 28.83 | 30.60 | 20.60 | 21.65 | 25.24 | 39.42 | 25.84 | 34.71 | 30.64 | 41.44 | 28.04 | 36.35 | 28.33 | 39.65 | 23.75 | 31.73 |

**Supporting Table 5.**
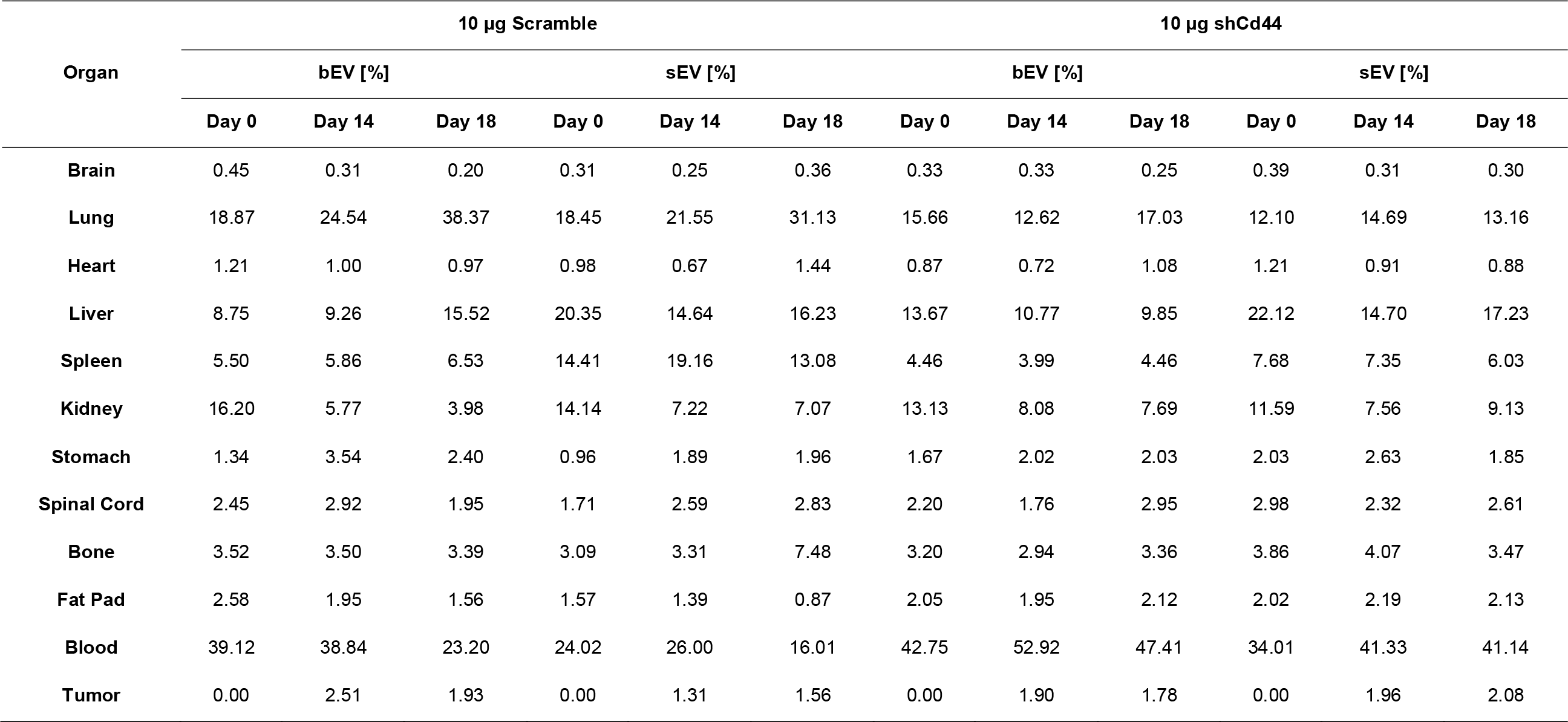
Organ biodistribution (%) of 4T1-PamGRET-Scramble, -shCd44 bEV and sEV at Days 0, 14, and 18 of EV treatment.

| Organ | 10 µg Scramble |  |  |  |  |  | 10 µg shCd44 |  |  |  |  |  |
| --- | --- | --- | --- | --- | --- | --- | --- | --- | --- | --- | --- | --- |
|  | bEV [%] |  |  | sEV [%] |  |  | bEV [%] |  |  | sEV [%] |  |  |
|  | Day 0 | Day 14 | Day 18 | Day 0 | Day 14 | Day 18 | Day 0 | Day 14 | Day 18 | Day 0 | Day 14 | Day 18 |
| **Brain** | 0.45 | 0.31 | 0.20 | 0.31 | 0.25 | 0.36 | 0.33 | 0.33 | 0.25 | 0.39 | 0.31 | 0.30 |
| **Lung** | 18.87 | 24.54 | 38.37 | 18.45 | 21.55 | 31.13 | 15.66 | 12.62 | 17.03 | 12.10 | 14.69 | 13.16 |
| **Heart** | 1.21 | 1.00 | 0.97 | 0.98 | 0.67 | 1.44 | 0.87 | 0.72 | 1.08 | 1.21 | 0.91 | 0.88 |
| **Liver** | 8.75 | 9.26 | 15.52 | 20.35 | 14.64 | 16.23 | 13.67 | 10.77 | 9.85 | 22.12 | 14.70 | 17.23 |
| **Spleen** | 5.50 | 5.86 | 6.53 | 14.41 | 19.16 | 13.08 | 4.46 | 3.99 | 4.46 | 7.68 | 7.35 | 6.03 |
| **Kidney** | 16.20 | 5.77 | 3.98 | 14.14 | 7.22 | 7.07 | 13.13 | 8.08 | 7.69 | 11.59 | 7.56 | 9.13 |
| **Stomach** | 1.34 | 3.54 | 2.40 | 0.96 | 1.89 | 1.96 | 1.67 | 2.02 | 2.03 | 2.03 | 2.63 | 1.85 |
| **Spinal Cord** | 2.45 | 2.92 | 1.95 | 1.71 | 2.59 | 2.83 | 2.20 | 1.76 | 2.95 | 2.98 | 2.32 | 2.61 |
| **Bone** | 3.52 | 3.50 | 3.39 | 3.09 | 3.31 | 7.48 | 3.20 | 2.94 | 3.36 | 3.86 | 4.07 | 3.47 |
| **Fat Pad** | 2.58 | 1.95 | 1.56 | 1.57 | 1.39 | 0.87 | 2.05 | 1.95 | 2.12 | 2.02 | 2.19 | 2.13 |
| **Blood** | 39.12 | 38.84 | 23.20 | 24.02 | 26.00 | 16.01 | 42.75 | 52.92 | 47.41 | 34.01 | 41.33 | 41.14 |
| **Tumor** | 0.00 | 2.51 | 1.93 | 0.00 | 1.31 | 1.56 | 0.00 | 1.90 | 1.78 | 0.00 | 1.96 | 2.08 |

**Supporting Table 6.**
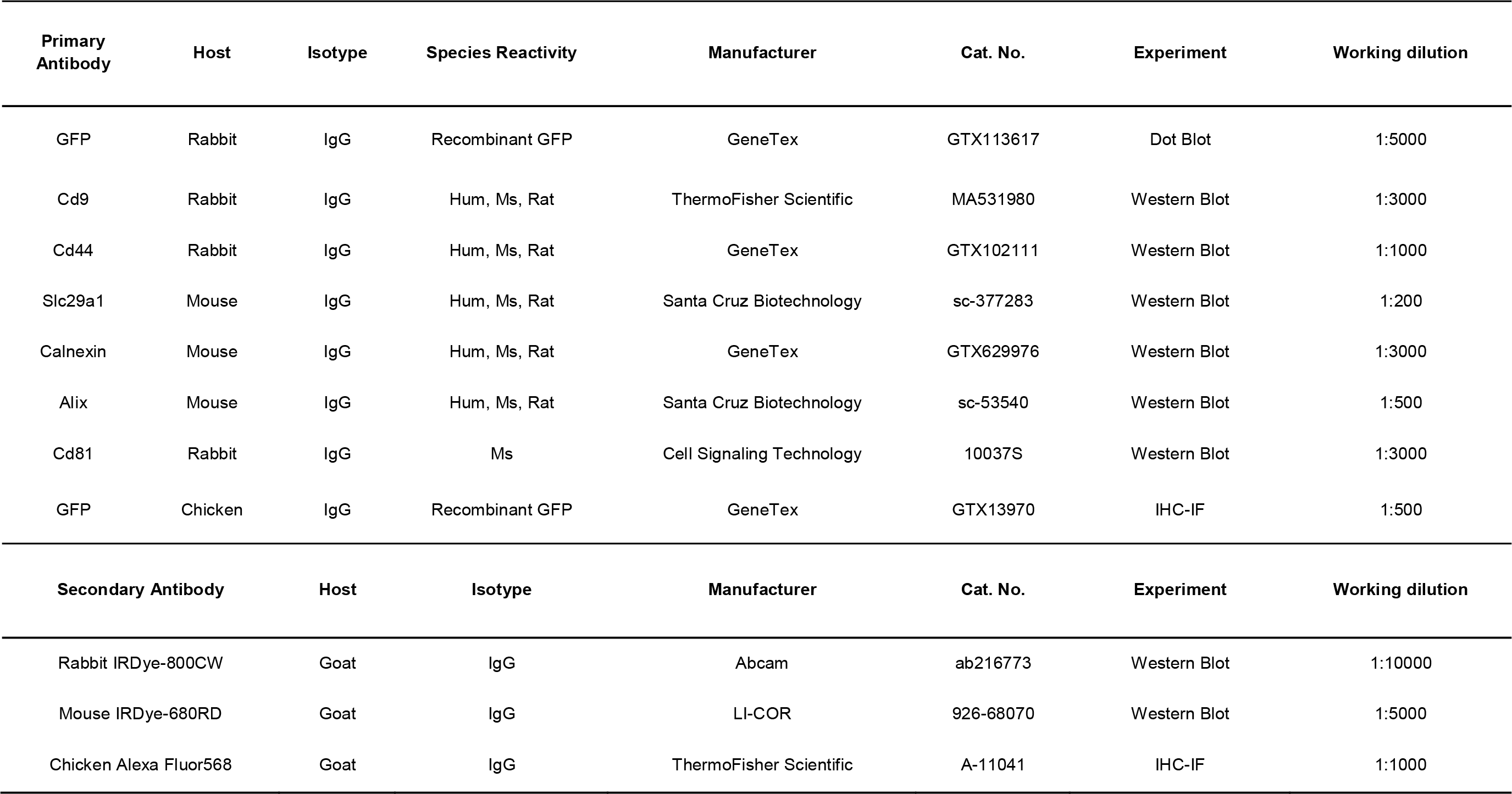
List of antibodies and working dilutions.

## References

1. B. T. Pan, K. Teng, C. Wu, M. Adam, R. M. Johnstone, The Journal of Cell Biology 1985, 101, 942 M. Tkach, C. Théry, Cell 2016, 164, 1226.

2. C. Théry, K. W. Witwer, E. Aikawa, M. J. Alcaraz, J. D. Anderson, R. Andriantsitohaina, A. Antoniou, T. Arab, F. Archer, G. K. Atkin-Smith, D. C. Ayre, J.-M. Bach, D. Bachurski, H. Baharvand, L. Balaj, S. Baldacchino, N. N. Bauer, A. A. Baxter, M. Bebawy, C. Beckham, A. Bedina Zavec, A. Benmoussa, A. C. Berardi, P. Bergese, E. Bielska, C. Blenkiron, S. Bobis-Wozowicz, E. Boilard, W. Boireau, A. Bongiovanni, F. E. Borràs, S. Bosch, C. M. Boulanger, X. Breakefield, A. M. Breglio, M. Á. Brennan, D. R. Brigstock, A. Brisson, M. L. Broekman, J. F. Bromberg, P. Bryl-Górecka, S. Buch, A. H. Buck, D. Burger, S. Busatto, D. Buschmann, B. Bussolati, E. I. Buzás, J. B. Byrd, G. Camussi, D. R. Carter, S. Caruso, L. W. Chamley, Y.-T. Chang, C. Chen, S. Chen, L. Cheng, A. R. Chin, A. Clayton, S. P. Clerici, A. Cocks, E. Cocucci, R. J. Coffey, A. Cordeiro-da- Silva, Y. Couch, F. A. Coumans, B. Coyle, R. Crescitelli, M. F. Criado, C. D’Souza- Schorey, S. Das, A. Datta Chaudhuri, P. de Candia, E. F. De Santana, O. De Wever, H. A. del Portillo, T. Demaret, S. Deville, A. Devitt, B. Dhondt, D. Di Vizio, L. C. Dieterich, V. Dolo, A. P. Dominguez Rubio, M. Dominici, M. R. Dourado, T. A. Driedonks, F. V. Duarte, H. M. Duncan, R. M. Eichenberger, K. Ekström, S. EL Andaloussi, C. Elie-Caille, U. Erdbrügger, J. M. Falcon-Perez, F. Fatima, J. E. Fish, M. Flores-Bellver, A. Försönits, A. Frelet-Barrand, F. Fricke, G. Fuhrmann, S. Gabrielsson, A. Gámez-Valero, C. Gardiner, K. Gärtner, R. Gaudin, Y. S. Gho, B. Giebel, C. Gilbert, M. Gimona, I. Giusti, D. C. Goberdhan, A. Görgens, S. M. Gorski, D. W. Greening, J. C. Gross, A. Gualerzi, G. N. Gupta, D. Gustafson, A. Handberg, R. A. Haraszti, P. Harrison, H. Hegyesi, A. Hendrix, A. F. Hill, F. H. Hochberg, K. F. Hoffmann, B. Holder, H. Holthofer, B. Hosseinkhani, G. Hu, Y. Huang, V. Huber, S. Hunt, A. G.-E. Ibrahim, T. Ikezu, J. M. Inal, M. Isin, A. Ivanova, H. K. Jackson, S. Jacobsen, S. M. Jay, M. Jayachandran, G. Jenster, L. Jiang, S. M. Johnson, J. C. Jones, A. Jong, T. Jovanovic-Talisman, S. Jung, R. Kalluri, S.-I. Kano, S. Kaur, Y. Kawamura, E. T. Keller, D. Khamari, E. Khomyakova, A. Khvorova, P. Kierulf, K. P. Kim, T. Kislinger, M. Klingeborn, D. J. Klinke, M. Kornek, M. M. Kosanović, Á. F. Kovács, E.-M. Krämer-Albers, S. Krasemann, M. Krause, I. V. Kurochkin, G. D. Kusuma, S. Kuypers, S. Laitinen, S. M. Langevin, L. R. Languino, J. Lannigan, C. Lässer, L. C. Laurent, G. Lavieu, E. Lázaro-Ibáñez, S. Le Lay, M.-S. Lee, Y. X. F. Lee, D. S. Lemos, M. Lenassi, A. Leszczynska, I. T. Li, K. Liao, S. F. Libregts, E. Ligeti, R. Lim, S. K. Lim, A. Linē, K. Linnemannstöns, A. Llorente, C. A. Lombard, M. J. Lorenowicz, Á. M. Lörincz, J. Lötvall, J. Lovett, M. C. Lowry, X. Loyer, Q. Lu, B. Lukomska, T. R. Lunavat, S. L. Maas, H. Malhi, A. Marcilla, J. Mariani, J. Mariscal, E. S. Martens-Uzunova, L. Martin-Jaular, M. C. Martinez, V. R. Martins, M. Mathieu, S. Mathivanan, M. Maugeri, L. K. McGinnis, M. J. McVey, D. G. Meckes, K. L. Meehan, I. Mertens, V. R. Minciacchi, A. Möller, M. Møller Jørgensen, A. Morales-Kastresana, J. Morhayim, F. Mullier, M. Muraca, L. Musante, V. Mussack, D. C. Muth, K. H. Myburgh, T. Najrana, M. Nawaz, I. Nazarenko, P. Nejsum, C. Neri, T. Neri, R. Nieuwland, L. Nimrichter, J. P. Nolan, E. N. Nolte-’t Hoen, N. Noren Hooten, L. O’Driscoll, T. O’Grady, A. O’Loghlen, T. Ochiya, M. Olivier, A. Ortiz, L. A. Ortiz, X. Osteikoetxea, O. Østergaard, M. Ostrowski, J. Park, D. M. Pegtel, H. Peinado, F. Perut, M. W. Pfaffl, D. G. Phinney, B. C. Pieters, R. C. Pink, D. S. Pisetsky, E. Pogge von Strandmann, I. Polakovicova, I. K. Poon, B. H. Powell, I. Prada, L. Pulliam, P. Quesenberry, A. Radeghieri, R. L. Raffai, S. Raimondo, J. Rak, M. I. Ramirez, G. Raposo, M. S. Rayyan, N. Regev-Rudzki, F. L. Ricklefs, P. D. Robbins, D. D. Roberts, S. C. Rodrigues, E. Rohde, S. Rome, K. M. Rouschop, A. Rughetti, A. E. Russell, P. Saá, S. Sahoo, E. Salas-Huenuleo, C. Sánchez, J. A. Saugstad, M. J. Saul, R. M. Schiffelers, R. Schneider, T. H. Schøyen, A. Scott, E. Shahaj, S. Sharma, O. Shatnyeva, F. Shekari, G. V. Shelke, A. K. Shetty, K. Shiba, P. R.-M. Siljander, A. M. Silva, A. Skowronek, O. L. Snyder, R. P. Soares, B. W. Sódar, C. Soekmadji, J. Sotillo, P. D. Stahl, W. Stoorvogel, S. L. Stott, E. F. Strasser, S. Swift, H. Tahara, M. Tewari, K. Timms, S. Tiwari, R. Tixeira, M. Tkach, W. S. Toh, R. Tomasini, A. C. Torrecilhas, J. P. Tosar, V. Toxavidis, L. Urbanelli, P. Vader, B. W. van Balkom, S. G. van der Grein, J. Van Deun, M. J. van Herwijnen, K. Van Keuren-Jensen, G. van Niel, M. E. van Royen, A. J. van Wijnen, M. H. Vasconcelos, I. J. Vechetti, T. D. Veit, L. J. Vella, É. Velot, F. J. Verweij, B. Vestad, J. L. Viñas, T. Visnovitz, K. V. Vukman, J. Wahlgren, D. C. Watson, M. H. Wauben, A. Weaver, J. P. Webber, V. Weber, A. M. Wehman, D. J. Weiss, J. A. Welsh, S. Wendt, A. M. Wheelock, Z. Wiener, L. Witte, J. Wolfram, A. Xagorari, P. Xander, J. Xu, X. Yan, M. Yáñez-Mó, H. Yin, Y. Yuana, V. Zappulli, J. Zarubova, V. Ž kas, J.-Y. Zhang, Z. Zhao, L. Zheng, A. R. Zheutlin, A. M. Zickler, P. Zimmermann, A. ė . Zivkovic, D. Zocco, E. K. Zuba-Surma, Journal of Extracellular Vesicles 2018, 7, 1535750.

3. K. Brennan, K. Martin, S. P. FitzGerald, J. O’Sullivan, Y. Wu, A. Blanco, C. Richardson, M. M. Mc Gee, Scientific Reports 2020, 10, 1039.

4. H. Zhang, D. Lyden, Nature Protocols 2019, 14, 1027.

5. J. W. Clancy, A. C. Boomgarden, C. D’Souza-Schorey, Nature Cell Biology 2021, 23, 1217.

6. C. Ciardiello, A. Leone, P. Lanuti, M. S. Roca, T. Moccia, V. R. Minciacchi, M. Minopoli, V. Gigantino, R. De Cecio, M. Rippa, L. Petti, F. Capone, C. Vitagliano, M. R. Milone, B. Pucci, R. Lombardi, F. Iannelli, E. Di Gennaro, F. Bruzzese, M. Marchisio, M. V. Carriero, D. Di Vizio, A. Budillon, Journal of Experimental & Clinical Cancer Research 2019, 38, 317.

7. F. Heitz, P. Harter, H. J. Lueck, A. Fissler-Eckhoff, F. Lorenz-Salehi, S. Scheil- Bertram, A. Traut, A. du Bois, Eur J Cancer 2009, 45, 2792 M. Riaz, M. T. M. van Jaarsveld, A. Hollestelle, W. J. C. Prager-van der Smissen, A. A. J. Heine, A. W. M. Boersma, J. Liu, J. Helmijr, B. Ozturk, M. Smid, E. A. Wiemer, J. A. Foekens, J. W. M. Martens, Breast Cancer Research 2013, 15, R33 A. Naderi, A. E. Teschendorff, N. L. Barbosa-Morais, S. E. Pinder, A. R. Green, D. G. Powe, J. F. Robertson, S. Aparicio, I. O. Ellis, J. D. Brenton, C. Caldas, Oncogene 2007, 26, 1507 X. Dai, H. Cheng, Z. Bai, J. Li, … Journal of Cancer 2017, 8, 3131.

8. O. Yersal, S. Barutca, World Journal of Clinical Oncology 2014, 5, 412.

9. L. Wang, B. Wang, H. Wen, J. Mao, Y. Ren, H. Yang, Oncol Rep 2020, 44, 407 S. Dutta, C. Warshall, C. Bandyopadhyay, D. Dutta, B. Chandran, PLoS One 2014, 9, e97580 S. Lucotti, C. M. Kenific, H. Zhang, D. Lyden, EMBO J 2022, 41, e109288 J. Peng, W. Wang, S. Hua, L. Liu, Breast Cancer (Auckl) 2018, 12, 1178223418767666.

10. S. T. Chuo, J. C. Chien, C. P. Lai, J Biomed Sci 2018, 25, 91 F. J. Verweij, L. Balaj, C. M. Boulanger, D. R. F. Carter, E. B. Compeer, G. D’Angelo, S. E. L. Andaloussi, J. G. Goetz, J. C. Gross, V. Hyenne, E.-M. Krämer-Albers, C. P. Lai, X. Loyer, A. Marki, S. Momma, E. N. M. N.-t. Hoen, D. M. Pegtel, H. Peinado, G. Raposo, K. Rilla, H. Tahara, C. Théry, M. E. v. Royen, R. E. Vandenbroucke, A. M. Wehman, K. Witwer, Z. Wu, R. Wubbolts, G. v. Niel, Nature Methods 2021, 18, 1013

11. M. Dehghani, S. M. Gulvin, J. Flax, T. R. Gaborski, Scientific Reports 2020, 1 A. Y.-T. Wu, Y.-C. Sung, Y. J. Chen, S. T. Y. Chou, V. Guo, J. C.-Y. Chien, J. J.-S. Ko, A. L. Yang, H. C. Huang, J. C. Chuang, S. Wu, M. R. Ho, M. Ericsson, W.-W. Lin, C. H. Y. Cheung, H. F. Juan, K. Ueda, Y. Chen, C. P.-K. Lai, Advanced Science 2020, 7, 2001467.

12. E. Lázaro-Ibáñez, F. N. Faruqu, A. F. Saleh, A. M. Silva, J. Tzu-Wen Wang, J. Rak, K. T. Al-Jamal, N. Dekker, ACS nano 2021, 15, 3212 N. G. Zhegalova, S. He, H. Zhou, D. M. Kim, M. Y. Berezin, Contrast Media Mol Imaging 2014, 9, 355.

13. G. F. Teare, P. K. Horan, S. E. Slezak, C. Smith, J. B. Hay, Cell Immunol 1991, 134, 157 M. Nowacki, L. Nazarewski, M. Pokrywczynska, T. Kloskowski, D. Tyloch, K. Pietkun, A. Jundzill, M. Rasmus, K. Warda, M. Gagat, A. Grzanka, M. Bodnar, A. Marszalek, M. Krawczyk, S. L. Habib, T. Drewa, Ann Transplant 2015, 20, 132 D. P. Kuffler, J Comp Neurol 1990, 302, 729.

14. T. Smyth, M. Kullberg, N. Malik, P. Smith-Jones, M. W. Graner, T. J. Anchordoquy, Journal of controlled release : official journal of the Controlled Release Society 2015, 199, 145 O. P. Wiklander, J. Z. Nordin, A. O’Loughlin, Y. Gustafsson, G. Corso, I. Mager, P. Vader, Y. Lee, H. Sork, Y. Seow, N. Heldring, L. Alvarez-Erviti, C. I. Smith, K. Le Blanc, P. Macchiarini, P. Jungebluth, M. J. Wood, S. E. Andaloussi, J Extracell Vesicles 2015, 4, 26316.

15. D. Gupta, A. Maria Zickler, S. EL Andaloussi, Advanced Drug Delivery Reviews 2021, 113961.

16. A. Hoshino, B. Costa-Silva, T.-L. Shen, G. Rodrigues, A. Hashimoto, M. T. Mark, H. Molina, S. Kohsaka, A. D. Giannatale, S. Ceder, S. Singh, C. Williams, N. Soplop, K. Uryu, L. Pharmer, T. King, L. Bojmar, A. E. Davies, Y. Ararso, T. Zhang, H. Zhang, J. Hernandez, J. M. Weiss, V. D. Dumont-Cole, K. Kramer, L. H. Wexler, A. Narendran, G. K. Schwartz, J. H. Healey, P. Sandstrom, K. J. Labori, E. H. Kure, P. M. Grandgenett, M. A. Hollingsworth, M. d. Sousa, S. Kaur, M. Jain, K. Mallya, S. K. Batra, W. R. Jarnagin, M. S. Brady, O. Fodstad, V. Muller, K. Pantel, A. J. Minn, M. J. Bissell, B. A. Garcia, Y. Kang, V. K. Rajasekhar, C. M. Ghajar, I. Matei, H. Peinado, J. Bromberg, D. Lyden, Nature 2015, 527, 329 H. Choi, Y. Choi, H. Y. Yim, A. Mirzaaghasi, J.-K. Yoo, C. Choi, Tissue Engineering and Regenerative Medicine 2021, 1.

17. A. Hoshino, B. Costa-Silva, T. L. Shen, G. Rodrigues, A. Hashimoto, M. Tesic Mark, H. Molina, S. Kohsaka, A. Di Giannatale, S. Ceder, S. Singh, C. Williams, N. Soplop, K. Uryu, L. Pharmer, T. King, L. Bojmar, A. E. Davies, Y. Ararso, T. Zhang, H. Zhang, J. Hernandez, J. M. Weiss, V. D. Dumont-Cole, K. Kramer, L. H. Wexler, A. Narendran, G. K. Schwartz, J. H. Healey, P. Sandstrom, K. J. Labori, E. H. Kure, P. M. Grandgenett, M. A. Hollingsworth, M. de Sousa, S. Kaur, M. Jain, K. Mallya, S. K. Batra, W. R. Jarnagin, M. S. Brady, O. Fodstad, V. Muller, K. Pantel, A. J. Minn, M. J. Bissell, B. A. Garcia, Y. Kang, V. K. Rajasekhar, C. M. Ghajar, I. Matei, H. Peinado, J. Bromberg, D. Lyden, Nature 2015, 527, 329 J. Mallegol, G. Van Niel, C. Lebreton, Y. Lepelletier, C. Candalh, C. Dugave, J. K. Heath, G. Raposo, N. Cerf-Bensussan, M. Heyman, Gastroenterology 2007, 132, 1866 S. Rana, S. Yue, D. Stadel, M. Zöller, The International Journal of Biochemistry & Cell Biology 2012 E. Segura, C. Guérin, N. Hogg, S. Amigorena, C. Théry, J Immunol 2007, 179, 1489.

18. M. Logozzi, A. De Milito, L. Lugini, M. Borghi, L. Calabrò, M. Spada, M. Perdicchio, M. L. Marino, C. Federici, E. Iessi, D. Brambilla, G. Venturi, F. Lozupone, M. Santinami, V. Huber, M. Maio, L. Rivoltini, S. Fais, PLoS ONE 2009, 4, e5219 M. P. Bebelman, M. J. Smit, D. M. Pegtel, S. R. Baglio, Pharmacol Ther 2018, 188, 1.

19. B. Bondhopadhyay, S. Sisodiya, F. A. Alzahrani, M. A. Bakhrebah, A. Chikara, V. Kasherwal, A. Khan, J. Rani, S. A. Dar, N. Akhter, P. Tanwar, U. Agrawal, S. Hussain, Cancers (Basel) 2021, 13.

20. A. Y.-T. Wu, Y.-C. Sung, Y.-J. Chen, S. T.-Y. Chou, V. Guo, J. C.-Y. Chien, J. J.-S. Ko, A. L. Yang, H.-C. Huang, J.-C. Chuang, S. Wu, M.-R. Ho, M. Ericsson, W.-W. Lin, C. H. Y. Cheung, H.-F. Juan, K. Ueda, Y. Chen, C. P.-K. Lai, Advanced Science 2020, 7, 2001467.

21. C. F. a. H. Bohren, D.R., in Absorption and Scattering of Light by Small Particles, 1998.

22. F. X. Schaub, M. S. Reza, C. A. Flaveny, W. Li, A. M. Musicant, S. Hoxha, M. Guo, J. L. Cleveland, A. L. Amelio, Cancer Research 2015, 75, 5023 N. Gaspar, J. R. Walker, G. Zambito, K. Marella-Panth, C. Lowik, T. A. Kirkland, L. Mezzanotte, Journal of Photochemistry & Photobiology, B: Biology 2021, 216, 112128.

23. N. Kastelowitz, H. Yin, ChemBioChem 2014, 15, 923.

24. J. C. Akers, V. Ramakrishnan, J. P. Nolan, E. Duggan, C.-C. Fu, F. H. Hochberg, C. C. Chen, B. S. Carter, PLOS ONE 2016, 11, e0149866 M. Wu, Y. Ouyang, Z. Wang, R. Zhang, P.-H. Huang, C. Chen, H. Li, P. Li, D. Quinn, M. Dao, S. Suresh, Y. Sadovsky, T. J. Huang, Proceedings of the National Academy of Sciences 2017, 114, 10584 M. Kanada, M. H. Bachmann, J. W. Hardy, D. O. Frimannson, L. Bronsart, A. Wang, M. D. Sylvester, T. L. Schmidt, R. L. Kaspar, M. J. Butte, A. C. Matin, C. H. Contag, Proceedings of the National Academy of Sciences 2015.

25. P. K. Wallace, J. D. T. Jr, J. L. Fisher, S. S. Wallace, M. S. Ernstoff, K. A. Muirhead, Cytometry Part A 2008, 73A, 1019

26. R. M. Abra, C. A. Hunt, Biochimica et Biophysica Acta (BBA) - Lipids and Lipid Metabolism 1981, 666, 493.

27. C. Y. Soo, Y. Song, Y. Zheng, E. C. Campbell, A. C. Riches, F. Gunn-Moore, S. J. Powis, Immunology 2012, Y. Xiao, W. Xu, Y. Komohara, Y. Fujiwara, H. Hirose, S. Futaki, T. Niidome, ACS Omega 2020, 5, 32744.

28. P. Gangadaran, X. J. Li, H. W. Lee, J. M. Oh, S. Kalimuthu, R. L. Rajendran, S. H. Son, S. H. Baek, T. D. Singh, L. Zhu, S. Y. Jeong, S. W. Lee, J. Lee, B. C. Ahn, Oncotarget 2017, 8, 109894.

29. J. Wu, P. Liu, J. L. Zhu, S. Maddukuri, M. A. Zern, Hepatology 1998, 27, 772 K. McEuen, J. Borlak, W. Tong, M. Chen, Int J Mol Sci 2017, 18.

30. S. W. Wen, J. Sceneay, L. G. Lima, C. S. F. Wong, M. Becker, S. Krumeich, R. J. Lobb, V. Castillo, K. N. Wong, S. Ellis, B. S. Parker, A. Möller, Cancer Research 2016, 76, 6816.

31. S. McFarlane, J. A. Coulter, P. Tibbits, A. O’Grady, C. McFarlane, N. Montgomery, A. Hill, H. O. McCarthy, L. S. Young, E. W. Kay, C. M. Isacke, D. J. Waugh, Oncotarget 2015, 6, 11465.

32. J. Cox, M. Mann, Nature Biotechnology 2008, 26, 1367.

33. Y. Perez-Riverol, J. Bai, C. Bandla, D. García-Seisdedos, S. Hewapathirana, S. Kamatchinathan, Deepti J. Kundu, A. Prakash, A. Frericks-Zipper, M. Eisenacher, M. Walzer, S. Wang, A. Brazma, Juan A. Vizcaíno, Nucleic Acids Research 2022, 50, D543.

34. H. Kalra, G. P. C. Drummen, S. Mathivanan, International Journal of Molecular Sciences 2016, 17.

